# Spatial Areas of Genotype Probability (SPAG): predicting the spatial distribution of adaptive genetic variants under future climatic conditions

**DOI:** 10.1101/2019.12.20.884114

**Authors:** Estelle Rochat, Stéphane Joost

**Affiliations:** Laboratory of Geographic Information Systems (LASIG), School of Architecture, Civil and Environmental Engineering (ENAC), Ecole Polytechnique Fédérale de Lausanne (EPFL), Lausanne, Switzerland

## Abstract

In a context of rapid global change, one of the key components for the survival of species is their genetic adaptive potential. Many methods have been developed to identify adaptive genetic variants, but few tools were made available to integrate this knowledge into conservation management. We present here the SPatial Areas of Genotype probability (SPAG), using genotype-environment logistic associations to map the probability of finding beneficial variants in a study area. We define a univariate model predicting the spatial distribution of a single genotype, and three multivariate models allowing the integration of several genotypes, potentially associated with various environmental variables. We then integrate climate change projections to map the corresponding future distribution of genotypes. The analysis of the mismatch between current and future SPAGs makes it possible to identify a) populations that are better adapted to the future climate through the presence of genetic variants able to cope with future conditions, and b) vulnerable populations where genotype(s) of interest are not frequent enough for the individuals to adapt to the future climate. We validate the SPAG approach using simulations and we use it to study the potential adaptation of 161 Moroccan and 382 European goats to the bioclimatic conditions. In Morocco, using whole genome sequence data, we identify seven genomic regions strongly associated with the precipitation seasonality (WorldClim database). The predicted shift in SPAGs under a strong climate change scenario for 2070 highlights goat populations likely to be threatened by the expected increase in precipitation variation in the future. In Europe, we find genomic regions associated with low precipitation, the shift in SPAGs highlighting vulnerable populations not adapted to the very dry conditions expected in 2070. The SPAG methodology is successfully validated using cross-validations and provides an efficient tool to take the adaptive potential into account in general conservation frameworks.

## Introduction

Climate change has altered the conditions of various organisms, causing an average warming of 0.2°C per decade over the past 30 years, as well as a sea level rise of more than 3 mm per year since the start of the century and an increase in the frequency of extreme weather events such as storms, droughts and floods (IPBES, 2019). These changes are likely to continue in the future (IPCC, 2014). When such important changes occur, many animal and plant species are confronted with a shift away from the favourable conditions necessary for their survival (IPBES, 2019). In order to avoid extinction under these conditions, they can either move to more favourable areas or adapt to their new environment (Hughes, 2000). Due to limitations in dispersal capacity, loss of favourable habitats and increased landscape fragmentation, the possibilities for dispersal to new areas are often limited (Opdam and Wascher, 2004; McGuire *et al.*, 2016). Population adaptation relies on phenotypic changes, which can be induced either by phenotypic plasticity or genetic evolution (Merilä and Hendry, 2014; Fox *et al.*, 2019). Phenotypic plasticity can allow species to rapidly evolve by changing their behaviour, physiology or morphology (Reed *et al.*, 2011; Fox *et al.*, 2019). However, it can also potentially lead to a fitness reduction (Duputié *et al.*, 2015) and, since it is not based on heritable genetic variations, it will not necessarily ensure the persistence of adaptation for the next generations. In order to preserve biodiversity, it is therefore crucial to promote the conservation of the genetic adaptive potential of populations (Hoffmann and Sgrò, 2011; Sgrò *et al.*, 2011; Nicotra *et al.*, 2015; Shafer *et al.*, 2015).

Conservation of this adaptive potential is also of major importance for livestock management in order to ensure herd persistence (Hoffmann, 2010; FAO, 2015). Indeed, many livestock populations, especially in developing country, are breed in pastoralist systems and live most of the time outdoors, confronted to difficult production conditions (e.g. heat stress, poor food resources and the presence of parasites and diseases), and without significant food or water supplies (FAO, 2015). These populations show signatures of local adaptation to their climatic conditions (McManus *et al.*, 2009, 2011; FAO, 2015; Bertolini *et al.*, 2018), to limited food resources (Silanikove, 2000) or to the presence of parasites (Noyes *et al.*, 2011; FAO, 2015; Vajana *et al.*, 2018). However, due to the increasing demand for food production, these local breeds currently tend to be replaced by high-producing commercial breeds imported from developed countries (Rischkowsky and Pilling, 2007; Hoffmann, 2010). This has led to a loss of genetic diversity, which threatens the adaptive potential of livestock species to environmental changes (FAO, 2015). In addition, imported breeds lack in the locally adapted genetic variants, which may reduce their fitness (FAO, 2015). It is therefore essential to highlight the adaptive potential of livestock species in order to encourage farmers to conserve local traditional breeds, and to carefully design cross-breeding, translocation or artificial selection (Scherf *et al.*, 2008; Allendorf *et al.*, 2010).

One of the essential components of the adaptive capacity of populations is genetic diversity (Allendorf and Leary, 1986). Since mutation rates are generally low, adaptation to rapid environmental changes largely depends on the amount of genetic variants already present in populations, i.e. standing genetic diversity (Orr and Unckless, 2008). With the recent increase in the availability of genetic data and the development of conservation genomics, various tools have been developed to integrate genetic diversity into conservation frameworks (Bonin *et al.*, 2007; Vandergast *et al.*, 2011; Thomassen *et al.*, 2011). However, conserving neutral variation in populations may not be sufficient to allow rapid adaptation to increasingly stressful conditions (Reed and Frankham, 2001), and it could be more valuable to specifically preserve adaptive variation, i.e. variation associated with a trait involved in fitness (Hoffmann and Willi, 2008; Sgrò *et al.*, 2011; Willoughby *et al.*, 2018) or to combine both approaches (Funk *et al.*, 2012; Pauls *et al.*, 2013). Increasing attention is currently being paid to this issue in conservation discussions (Funk *et al.*, 2019; Hoelzel *et al.*, 2019; Mable, 2019).

Several methods have been developed to identify signatures of local adaptation, based on various assumptions and with different limitations and advantages (Schoville et al., 2012; Joost et al., 2013; Vitti et al., 2013; Hoban et al., 2016). The results have notably been used to establish prediction of future habitat range of species facing climate change (Hällfors et al., 2016; Ikeda et al., 2017; Garzón et al., 2019; Razgour et al., 2019). However, there is currently a need to integrate this knowledge in order to predict the distribution of locally adapted genetic variants along environmental gradients, and to project the probability of finding them in un-sampled areas or under future climatic conditions (Bay et al., 2017). Nevertheless, very few studies have addressed this issue. Fournier *et al.* (2011) identified locally adapted SNP alleles and then used the Maxent species distribution model (Phillips et al., 2006) to estimate their spatial distribution. Fitzpatrick and Keller (2015) proposed another approach based on two community-level modelling methods (Generalised Dissimilarity Modelling and Gradient Forest). They used it to map the current and future spatial distribution of several adaptive variants and to assess the “genetic offset” of populations under climate change as a function of the mismatch between the current distributions and future predictions. However, practical applications of these methods remain limited and there is still a need to develop new tools to ease the integration of the adaptive potential into conservation practices, especially by considering several loci, potentially non-independent and adapted to different environmental conditions.

We propose here a novel approach to predict genotype frequencies and map SPatial Areas of Genotypes Probabilities (SPAG) based on logistic genotype-environment associations (Joost *et* al., 2007) and conditional probability theory. SPAGs can be used to a) predict the probability of presence of one or many locally adapted genetic variants in non-sampled areas b) identify areas where there is a greater probability of finding individuals better adapted to future climatic conditions, c) identify vulnerable populations that may be threatened by climate change and d) integrate the results into conservation frameworks by means of an easy combination with other georeferenced layers. The concept of applying logistic regressions on an environmental layer to predict the probability of presence of a genotype had been sketched out several years ago (Joost 2006; page 138). Here, we formalise this concept and extend it to multivariate models. We introduce the theoretical bases of SPAGs and validate the approach with a simulated dataset. We then present an application of our approach to two case studies in order to analyse the local adaptive potential of Moroccan and European goat populations.

## Material and Methods

### SPAG’s approach

#### Logistic regressions (SAM)

The Spatial Analysis Method (SAM, Joost *et al.*, 2007) can be used to detect genotypes that are strongly associated with an environmental variable and are therefore potential adaptive variants (strictly additive). This method assumes a linear response of the genotype to the environmental variable and uses logistic regressions (Formula 1) to assess the probability of presence of a genotype *G1* as a function of an environmental variable *(x1)*,

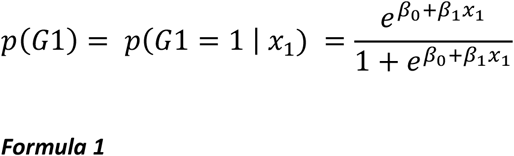

where β0 and β1 are the parameters of the regression to be fitted. Independent univariate logistic regressions can be computed between each genotype and each environmental predictor and significant associations can be identified using statistical tests. Joost *et al.* (2007) suggested the combined use of a likelihood ratio (G score) and a Wald test. The likelihood ratio (G) compares the likelihood of a model with the likelihood of a null model without the environmental variable of interest. The null hypothesis is that the model considered is no better than the null model, i.e. the environmental variable considered does not help in predicting the probability of presence of the genotype. The Wald test is a common statistic used to estimate whether a parameter is equal to a given value. In our case, it is used to reject the null hypothesis that the β parameter associated with an environmental variable is equal to 0, which would also indicate that it has no effect on the probability of finding the genotype. The SAM approach has been implemented in the Samβada software and validated against other methods for identifying signatures of natural selection (Stucki *et al.*, 2017).

#### Univariate SPAG

Once the genotype(s) involved into significant associations with the environmental variables have been identified, Formula 1 enables the estimation of the probability of presence of a genotype for any value of an environmental variable (*x1*). We consequently used it to estimate and delimit on a map the probability of presence of a genotype over the whole region of interest (Joost, 2006; Rochat *et al.*, 2016). We named such a delimited surface univariate Spatial Area of Genotype Probability (SPAG).

As more than one adaptive locus are usually identified, we also developed multivariate models to compute a single map showing the probability of presence of multiple genotypes. Three different multivariate models were developed to date: the Intersection, Union and K-Percentage.

#### Intersection SPAG (I-SPAG)

The **Intersection** model (I-SPAG) is used to compute the probability that the variants of interest are all simultaneously present. Following the theory of conditional probability (Kolmogorov, 1956), the probability of simultaneous presence of *n* adaptive genotypes *Gi, i=1:n* can be computed using Formula 2:

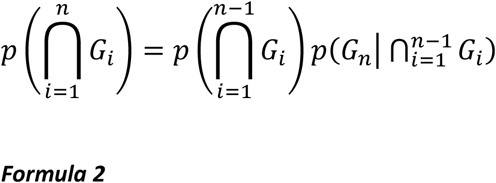

where 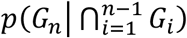 is a conditional probability that can be estimated using a logistic regression where 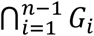 is integrated as a covariate (Formula 3).

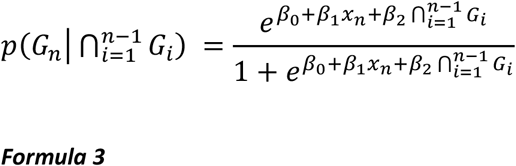

However, as we would like to use this model to predict the probability of presence of the genotypes for any point of the region of interest, i.e. also where *Gi* values are unknown, we suggested to estimate 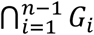 by 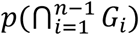.

Using the associative property of the intersection operator, the intersection of *n* genotypes can be computed by starting with the univariate model *p(G1)*, which is used as a covariate to compute *p(G1 ∩ G2)*, itself used to compute *p(G3 ∩ (G1 ∩ G2))*, etc.. Formula 3 can thus be implemented with a recursive model based on the univariate formula in which covariates are added (see Supp. File 1 for more details). We implemented this model as an *R* function available following the link given in the section “code availability” at the end of the manuscript.

#### Union SPAG (U-SPAG)

The **union** model (U-SPAG) is used to compute the probability of finding at least one of the adaptive genotypes of interest. We implemented it with the inclusion-exclusion principle (e.g. for two genotypes: p(G1 ∪ G2) = p(G1) + p(G2) – p(G1 ∩ G2)). We implemented the generalised formula for *n* adaptive genotypes (Formula 4), as an *R* function based on the intersection model previously described (see code availability and Suppl. File 1).

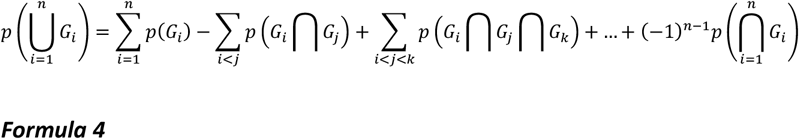

#### K-Percentage SPAG (K-SPAG)

Finally, we developed a **K-percentage** model (K-SPAG) to estimate the probability that an individual carries K% of *n* adaptive genotypes. This probability can be computed by combining formulas from the union and intersection models (Formula 5, explained in more detail in Supp. File 1). Again, this formula was implemented as an *R* function (see code availability).

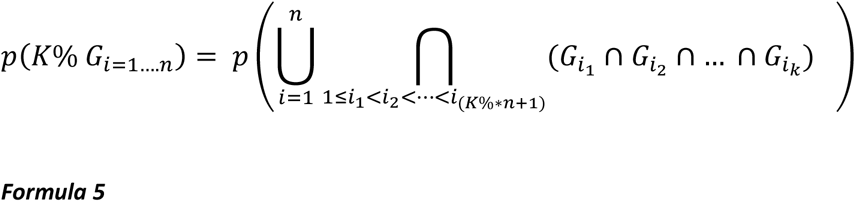

Note that all multivariate models allow the integration of adaptive genotypes associated with various environmental variables since the environmental variable *xi* used to compute *p(Gi)* can be different for each *i*.

### Simulation study

In order to test the SPAG’s approach, we first computed a simulated dataset using the individual-based population genetics model software CDPOP 1.3 (Landguth and Cushman, 2010; Landguth *et al.*, 2020). We simulated individual genetic exchanges and natural selection across 300 non-overlapping generations among 200 individuals randomly located in a 500×500 gridded landscape. For the breeding parameters, we considered a sexual reproduction, with random mating, both male and female with replacement, no selfing, no philopatry, no multiple paternity, equal sex ratio and each mated pair producing three offspring. The movement of the individuals was linearly restricted as a function of the Euclidean distance, with a maximum dispersal corresponding to 25% of the entire landscape. We simulated 50 diallelic loci, with three loci under selection (L0, L1 and L2). The selection was implemented using three 500×500 raster gradients, the first from north to south (X0), the second from east to west (X1) and the third from northwest to southeast (X2) (see Figure1A). We set the average effects bL0A0A0=10 and bL0A1A1=−10 for the locus L0 with the environmental variable X0, which indicates that the genotype A0A0 from locus L0 will be favoured in the South (where X0=1), whereas A1A1 will be favoured in the North (where X0=−1). We set similar effects for the locus L1 with the environmental variable X1 and L2 with X2. All other beta effects were set to 0, indicating no influence of the environmental variable to the distribution of genotypes. All genotypes were randomly initialised at the beginning of the simulations. The exact list of simulation parameters used is provided in Supp. File 2.

**Figure 1.**
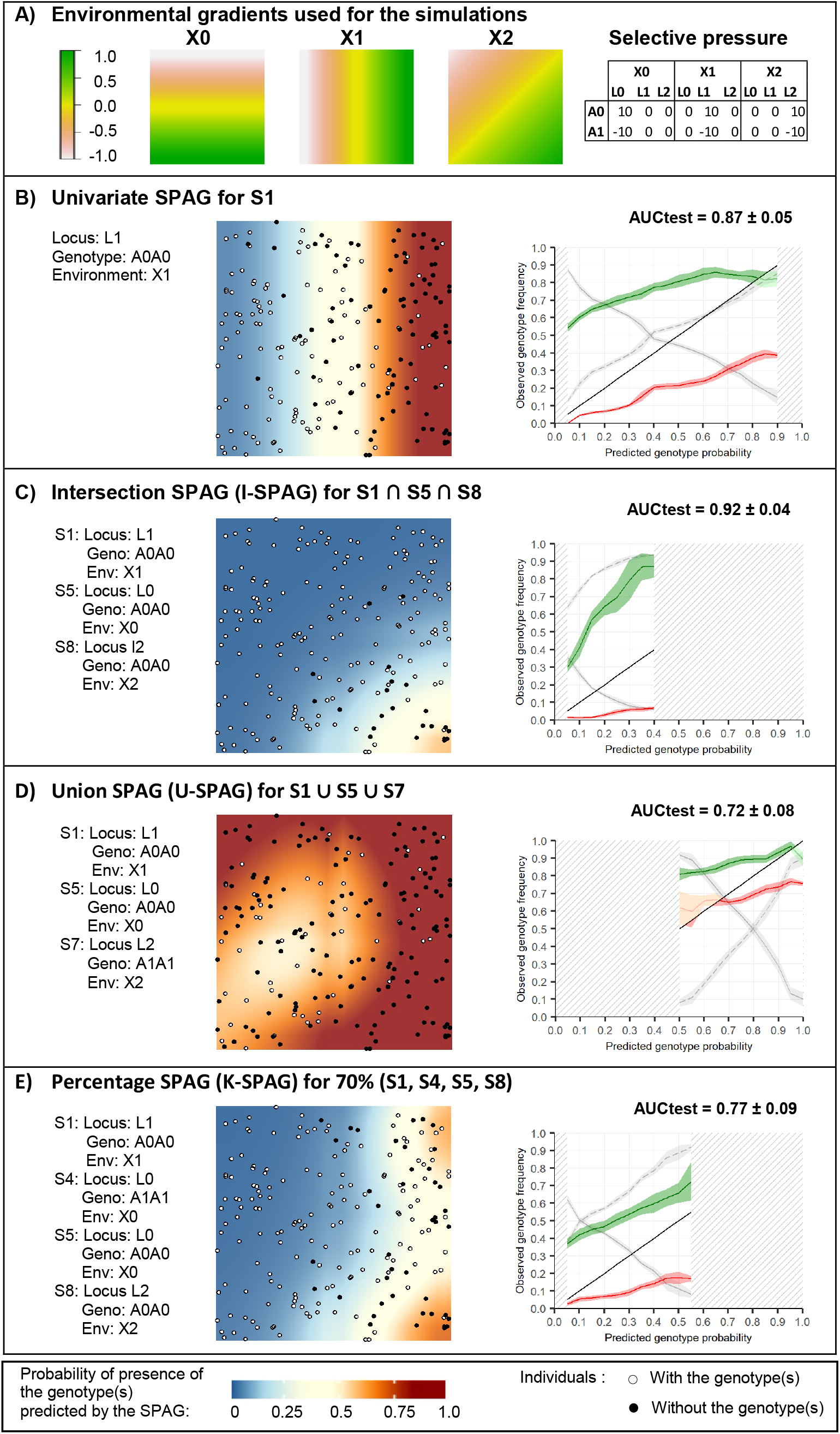
SPAG – Simulated dataset. Univariate and Multivariate Spatial Areas of Genotypes Probability for the simulated dataset. The identifiers of the presented models (S1, S4, S5, S7, S8) refers to Table 1. Please refer to Box 1 to interpret the validation graphs shown on the right of each map.

Univariate logistic regressions were then applied to the genetic data of individuals at the 300^th^ generation in order to identify the most significant associations, which should highlight loci under selection. Univariate and multivariate SPAGs were then applied to estimate the probability to find f finding these genotypes across the simulated landscape. The results were validated using a cross-validation procedure presented below.

### Moroccan and European goats

#### Genetic data

Two genetic datasets characterising goats (Capra hircus) were used for the analyses presented here. The first one was produced in the context of the NEXTGEN project (Alberto *et al.*, 2018) and the second was collected by the ADAPTMAP consortium (Stella *et al.*, 2018; http://www.goatadaptmap.org/).

The NEXTGEN project produced whole genome sequences data for 161 Moroccan goats from 6 different local breeds. Since goat production system in Morocco is mainly free-range, these goats are living from 8 to 12 months outdoors (Boujenane, 2005), and are confronted to contrasting environmental conditions, from the Sahara desert to the Atlas Mountains (see Figure A in Suppl. File 3). The goats were sampled in 161 farms chosen such to be representative of the range of environmental conditions observed in Morocco (Stucki, 2014). The sequencing method is described by Benjelloun *et al.* (2015) and allows to genotype 31.8 M of SNPS mapped to the goat’s reference genome CHIR v1.0 (Dong *et al*., 2013).

The ADAPTMAP consortium gathered genetic data for 4’563 goats from 144 breeds, sequenced worldwide with the CaprineSNP50 BeadChip and mapped on the most recent goat reference genome ARS1 (Bickhart *et al.*, 2017). The goats were georeferenced to the place where they have been sampled. We used here a subset of these data, constituted of individuals from Switzerland, North of Italy and France. This represented 458 individuals distributed in 196 locations, with 1 to 39 individuals per site. In order to avoid overweighting of some locations, we selected a maximum of five individuals per sampling site. These five individuals were chosen as the subset showing the highest Nei’s genetic distances, computed with the function *dist.genpop* from the package *adegenet* (Jombart, 2008) in the *R* environment (R Development Core Team, 2008). The resulting dataset contains 382 individuals from 196 locations and 11 different breeds (see Figure B in Supp. File 3).

Both genetic datasets were filtered such as to keep only autosomal, bi-allelic SNPs, with a maximum missingness per individuals and per site of 0.05 and a maximum major genotype frequency of 0.9. The final datasets contain 8,497,971 SNPs for the Moroccan goats and 46,294 SNPs for the European ones.

#### Environmental data

The climatic conditions of the sampling locations were characterised using the 19 bioclimatic variables (Supp. File 4) from the WorldClim database (https://www.worldclim.org/), representative of the period 1960-1990 (Hijmans *et al.*, 2005). Each variable was retrieved as a raster layer with a spatial resolution of 30 arc-seconds (approx. 1km2) and values were extracted for all sampling locations using the *extract* function from the R-package *raster* (Hijmans and van Etten, 2012). In order to get similar ranges of values for all bioclimatic variables, which makes it easier to compare the subsequently derived models, all variables were standardised for each dataset, by subtracting the mean and dividing by the standard deviation. Some of the bioclimatic variables are highly correlated. However, we choose to keep all of them to be able to identify *a posteriori* which variable had the strongest effect. Since no models computed involved more than one environmental variable simultaneously, this collinearity will not impact the results.

#### Population Structure

The genetic population structure was estimated with a Principal Component Analysis (Price *et al.*, 2006; Reich *et al.*, 2008) computed with the function *snpgdsPCA* from the *SNPRelate* R-package (Zheng *et al.*, 2012). In order to avoid a strong influence of SNP clusters on this analysis, we used here a pruned set of SNPs that are in approximate linkage equilibrium with each other. The pruning was performed with the function *snpgdsLDpruning* from the *SNPRelate* package, with a threshold D’=0.2. The resulting datasets contain 59,224 SNPs for the Moroccan goats and 14,571 SNPs for the European ones.

#### Logistic regressions and SPAGs

For the two datasets, logistic models were computed for each genotype with the 19 bioclimatic variables. The statistical significance of the model was assessed using Wald test and log-likelihood ratio (G), both corrected for the false-discovery rate due to multiple comparisons using the procedure proposed by Benjamini and Hochberg (1995), under an expected false discovery rate (FDR) of 0.05 (i.e. 5% of the results expected to be false positives).

In order to lower the number of false positive resulting from demographic processes instead of natural selection (Li *et al.*, 2012), logistic models were computed with the addition of covariates corresponding to the coordinates of individuals on the significant components of the PCA. The significance of the models with population covariates was assessed using a Wald test and a log likelihood ratio which compares the model with environment and covariates to the model with covariates only. An association was considered as significant if both the models without covariates and with population covariates were significant.

Finally, to identify potential functions of the SNPs involved into the significant associations, we used the NCBI Genome Data Viewer (https://www.ncbi.nlm.nih.gov/genome/gdv/browser/genome/?id=GCF_000317765.1) to search for the presence of annotated genes in the genomic region of 10kbp surrounding the SNPs of interest. All analyses were computed using a combination of the Samβada software (Stucki *et al.*, 2017) and a custom R-script based on the *glm* function.

### Validation procedure

For the simulated data as well as for the two goat datasets, SPAGs were validated with a cross-validation procedure, using 25% of the individuals to compute the SPAG (i.e. 50 individuals for the simulated datasets, 41 individuals for the Moroccan goats and 96 for the European) and the remaining 75% to test it. Training individuals were selected such to represent the entire range of values of the environmental variable under study (see Supp. File 1 for the exact procedure). The model was validated using the Area Under the Receiver Operating Curve computed with the testing dataset (AUCtest, (Fielding and Bell, 1997)) and a custom validation graph presented in Box 1. The cross-validation procedure was repeated 10 times.

#### Box 1

##### Validation Procedure

SPAGs indicate the probability of finding one or more genotypes of interest in a territory (panel below). For a given threshold value (e.g. th=0.6), we can thus use the SPAG to delimit the area where the probability of finding the genotype(s) of interest is predicted to be greater or equal to this threshold (e.g. probability>=0.6). If the SPAG is valid, the frequency of the genotype(s) observed among the testing individuals located within the thresholded SPAG should effectively be greater or equal to the threshold value, whereas it should be less outside.

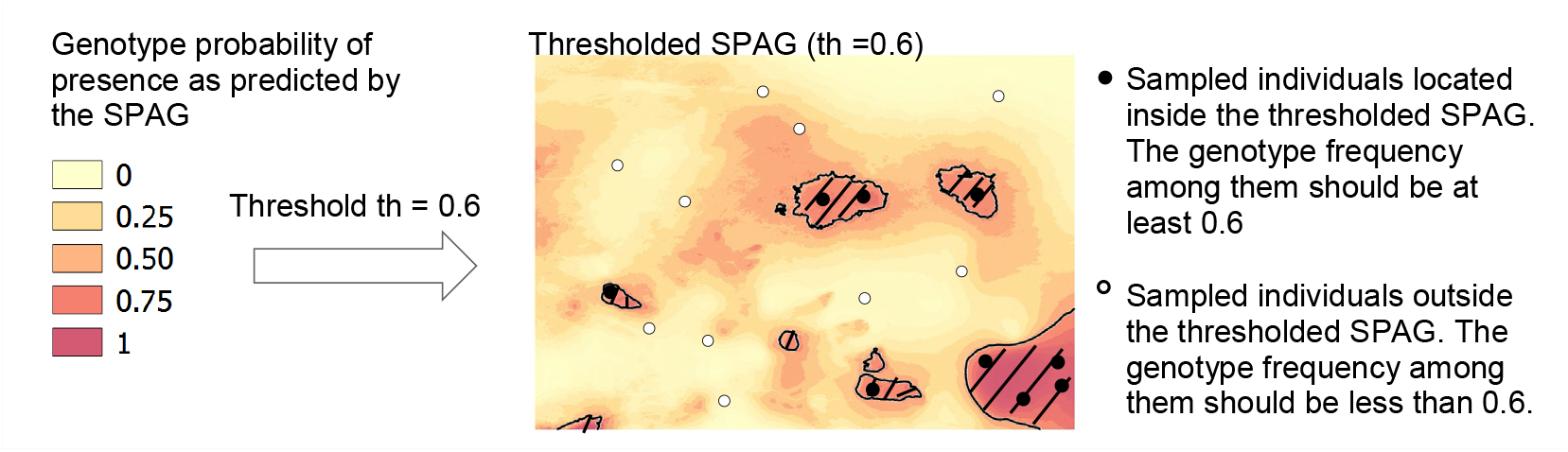

To validate the SPAGs, we thus calculated the observed genotype frequencies inside and outside the thresholded SPAGs for each threshold value between 0 and 1 (with a step of 0.1) and we presented the results on a graph (panel below). The green line indicates the genotype frequency observed inside the thresholded SPAG, whereas the red line shows the genotype frequency observed outside it. A black line indicates the limit case where the observed genotype frequency is equal to the threshold value. The SPAG is thus validated if the green line remains above the black line and the red line remains below it. The green and red areas around the lines indicate the 95% confidence intervals for each line, computed on the basis of the 10 cross-validation runs. We also presented on the graph the percentage of individuals located within the thresholded SPAG (grey line) and outside it (dotted grey line). The hatched grey areas indicate ranges of testing values where there was less than 5 individuals remaining inside or outside the thresholded SPAG, which was therefore considered not to be usable for the validation.

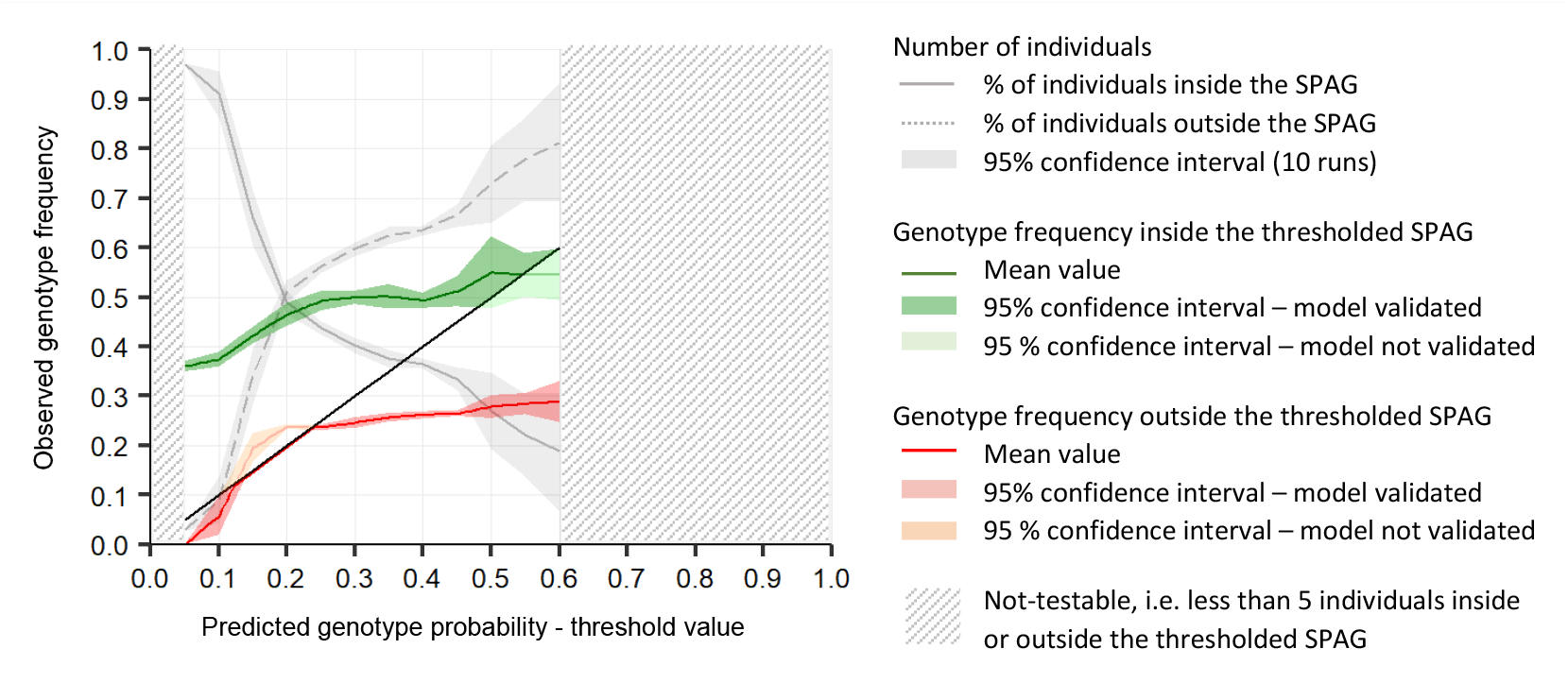

*For a threshold value th=0.4, the SPAG is validated since 50% of the testing individuals located within the area SPAG≥0.4 carry the genotype of interest, whereas only 26% carry it ouside (SPAG<0.4). Inversely, the model is not validated for th=0.6, since only 54% of individuals carry the genotype of interest whithin the area where the SPAG predicted a probability of at least 0.6 (SPAG ≥0.6). The model is also not valid for a value th=0.2 since 23% of individuals carry the genotype of interest in the area SPAG<0.2.*

### Projections under climate change

We use the two goat datasets to present an application of the SPAG for predicting the distribution of adaptive genotype(s) under climate change. In order to predict the genotype frequency optimal for future conditions, we retrieved Worldclim data for the year 2070, corresponding to a strong climate change scenario from the Max Planck Institute Earth System Model (MPI-ESM-LR) (Giorgetta *et al.*, 2013) with a Representative Concentration Pathway equals to 8.5 (RCP 8.5). We then assume that the optimal genotype frequency for future conditions should be close to the genotype frequency currently observed in areas with climatic conditions resembling the future ones. We thus applied the current parameters of the logistic regressions on the future environmental variables in order to derive the future SPAGs for the genotypes of interest. We then study the shift between the current and future SPAGs to identify vulnerable populations for which specific genotype frequencies should be much higher so that individuals can adapt to the future conditions.

## Results

### Simulated results

The 10 most significant genotype-environment associations obtained with the simulated datasets ranked on the basis of the likelihood ratio (G) are presented in Table 1. We observe that the three loci simulated as under selection (L0, L1, L2) are coherently identified as the most significantly associated with the environmental variables under study.

**Table 1.** Most significant models – Simulated datasets. 10 most ignificant models obtained for the analysis of the simulated datasets, ranked based on the likelihood ratio. Geno = Genotype, Env=environmental predictor, pG=p-value associated with the G score, pW=p-value associated with the Wald score, β0 and β1 are the parameters of the logistic regression.

| ID | Marker | Locus | Geno. | Env. | G score | pG | pW | $\beta_0$ | $\beta_1$ |
| --- | --- | --- | --- | --- | --- | --- | --- | --- | --- |
| S1 | L1A0A0 | L1 | A0A0 | X1 | 92.30 | 7.44E-22 | 9.41E-14 | -0.46 | 2.65 |
| S2 | L1A1A1 | L1 | A1A1 | X1 | 90.49 | 1.86E-21 | 1.51E-12 | -1.24 | -2.94 |
| S3 | L2A1A1 | L2 | A1A1 | X2 | 68.63 | 1.19E-16 | 9.02E-11 | -1.59 | -4.37 |
| S4 | L0A1A1 | L0 | A1A1 | X0 | 65.69 | 5.29E-16 | 4.04E-11 | -0.66 | -2.35 |
| S5 | L0A0A0 | L0 | A0A0 | X0 | 64.95 | 7.69E-16 | 6.92E-11 | -1.58 | 2.61 |
| S6 | L2A0A0 | L2 | A0A0 | X1 | 63.86 | 1.34E-15 | 1.37E-11 | -0.55 | 2.04 |
| S7 | L2A1A1 | L2 | A1A1 | X1 | 61.04 | 5.59E-15 | 4.12E-10 | -1.56 | -2.40 |
| S8 | L2A0A0 | L2 | A0A0 | X2 | 60.94 | 5.88E-15 | 2.52E-10 | -0.54 | 3.45 |
| S9 | L0A0A0 | L0 | A0A0 | X2 | 43.43 | 4.39E-11 | 2.81E-08 | -1.41 | 3.11 |
| S10 | L42A0A0 | L42 | A0A0 | X0 | 42.44 | 7.30E-11 | 5.93E-09 | 0.06 | 1.67 |

Figure 1B presents the univariate SPAG for the model S1 (Locus L1, genotype A0A0 associated with the environmental variable X1). Since this locus was simulated as under selection with the east-west gradient X1, the resulting SPAG coherently shows a similar gradient. The results are validated by the validation graph for almost the entire range of probability values, except close to 0.8, where the SPAG slightly overestimate the probability of presence of the genotype (the green line falls below the black one, i.e. the observed genotype frequency is lower than what predicted by the SPAG). Figure 1C shows the intersection SPAG for the genotype A0A0 of the three loci under selection. It indicates a very low probability of finding them all simultaneously, except in the South-East of the simulated area (where all environmental gradients considered had values close to 1). Figure 1D presents a union-SPAG of three genotypes. It shows that the probability of finding at least one of them is higher than 0.5 in most of the area. The validation graph indicates that the model tends to underestimate the probability of finding the genotypes for threshold values below 0.6 (the red line is above the black one, i.e. the genotype frequency outside the thresholded SPAG is higher than what predicted). Finally, Figure 1E depicts the probability of finding 70% of 4 genotypes of interest, i.e. at least 3 of them. This probability is low, except in the South-East and North-West part. Again, the validation graph indicates a good power of the SPAG to predict the probability of finding a set of genotypes of interest.

### Moroccan goats

#### Population structure

For the Moroccan dataset, the cumulated variance explained by the 10 first PCA components on the SNP markers represents only 8.1% of the total variance and the increase in variance explained is almost proportional to the number of components, which highlights that there is no clear sub-structure. We therefore do not include any population structure on the subsequent analysis and computed only logistic regressions without any covariates.

#### Logistic regressions

More than 483 million logistic association models were computed. After correction for false discovery rate with a significant threshold of 5%, no model is significant according to the Wald score, but seven models are significant according to the G score (Supp. File 5). Among them, three models were strongly associated with the precipitation seasonality (bio15), which is a measure of the variation of monthly precipitation over the year. Following this initial result, we investigated in more details the adaptation to this bioclimatic variable. When considering only the associations involving bio15 (25,447,348 models), 78 models are significant after FDR-correction of G score, with a significant threshold of 5% (Supp. File 5). The SNPs involved in these models are located on seven different genomic regions (Table 2), corresponding to four annotated genes (DSG4, CDH2, KCTD1 and WRN) on the reference genome CHIR 1.0.

**Table 2.** Significant models – Moroccan datasets – Bio15. Significant models obtained for the analysis of Moroccan datasets with precipitation seasonality (bio15) after FDR correction. Chr=Chromosome, Start=Start in base pairs of the region identified as under selection, End=End in base pairs of the region, Peak SNP = SNP of the most significant model in that region, Geno = corresponding SNP Genotype, GF=corresponding Genotype Frequency, β0 and β1 = parameters of the logistic regression, G=G score (Log Likelihood ratio), qG=corresponding p-value corrected for FDR, Genes = Annotated genes on the genomic region.

| ID | Chr | Start (BP) | End (BP) | Peak (BP) | Geno | GF | G | qG | $\beta_0$ | $\beta_1$ | Genes |
| --- | --- | --- | --- | --- | --- | --- | --- | --- | --- | --- | --- |
| M1 | 6 | 12'174'332 | 12'298'321 | 12'276'168 | AA | 21.74 | 38.74 | 0.004 | -1.80 | 1.51 | - |
| M2 | 13 | 43'436'394 | 43'438'732 | 43'436'394 | GG | 10.56 | 29.02 | 0.042 | -3.14 | 1.80 | - |
| M3 | 24 | 19'436'980 | 19'436'980 | 19'436'980 | CC | 76.40 | 34.75 | 0.008 | 1.55 | 1.29 | - |
| M4 | 24 | 25'852'900 | 25'860'754 | 25'860'754 | AG | 38.51 | 34.75 | 0.008 | -0.29 | -1.07 | DSG4 |
| M5 | 24 | 28'799'029 | 28'833'762 | 28'833'253 | TT | 12.42 | 27.87 | 0.046 | -2.72 | 1.58 | CDH2 |
| M6 | 24 | 30'566'869 | 30'584'692 | 30'566'869 | TT | 2.48 | 27.99 | 0.046 | -25.69 | -15.44 | KCTD1 |
| M7 | 27 | 25'930'079 | 25'933'133 | 25'930'079 | GG | 78.88 | 32.88 | 0.012 | 1.76 | -1.35 | WRN |

#### Spatial Areas of Genotype Probability

Figure2A shows the univariate SPAG for the genotype of model M1 presented in Table 2 (see Supp. File 7 for the other univariate SPAGs). The predicted probability of presence of the genotype is the highest in the extreme southwest of the country, near the Sahara desert. In this region, the variations of precipitation are the greatest (the standard deviation of monthly precipitation is more than 100% of the mean of monthly precipitation) and all goats carry the genotype of interest. In coastal areas, the predicted probability of finding the genotype is close to 0.5. In this area, the variations of precipitations are also high (more than 70%) and some of the sampled goats carry the genotype, while others do not. Finally, in the Atlas Mountains and in the northeast of the country, the probability of finding the genotype of model M1 is much lower (<0.2 in most areas). In these regions, variations of precipitation are less important (35-50%) and most of the goats sampled do not carry the genotype.

Two other markers positively correlated with bio15 were highlighted by models M3 and M5 (Table 2). Nevertheless, the I-SPAG presented in Figure 2B indicates that their simultaneous presence is very unlikely (probability <0.1 for most of the territory). On the contrary, the probability of finding at least one of the genotypes from the three models M1, M3 and M5, all positively associated with the coefficient of precipitation, is very high in many parts of the territory (U-SPAG, Figure 2C). Finally the K-SPAG presented in Figure 1D shows the probability that goats carry at least 50% of the four variants positively associated with the coefficient of precipitation (M1, M2, M3, M5), i.e. the probability to find at least two of them. This map is the most contrasted, showing a very high probability of presence near the coast and the Sahara desert (> 0.9) and a very low probability (<0.2) in the centre and northeast of the country. For all these multivariate cases, the mean AUC value for the testing dataset over the 10 runs is greater than 0.8. In addition, the validation graphs indicate that the SPAGs computed with 25% of the individuals generally enable a correct estimate of the probability of finding the genotype(s) of interest in the 75% remaining individuals. Only the U-SPAG (Figure 2C) tends to slightly overestimate the probability of presence since we observe a higher presence of the genotypes in the individuals located outside the thresholded SPAG as compare to what predicted by the SPAG (the red line is above the black line).

**Figure 2.**
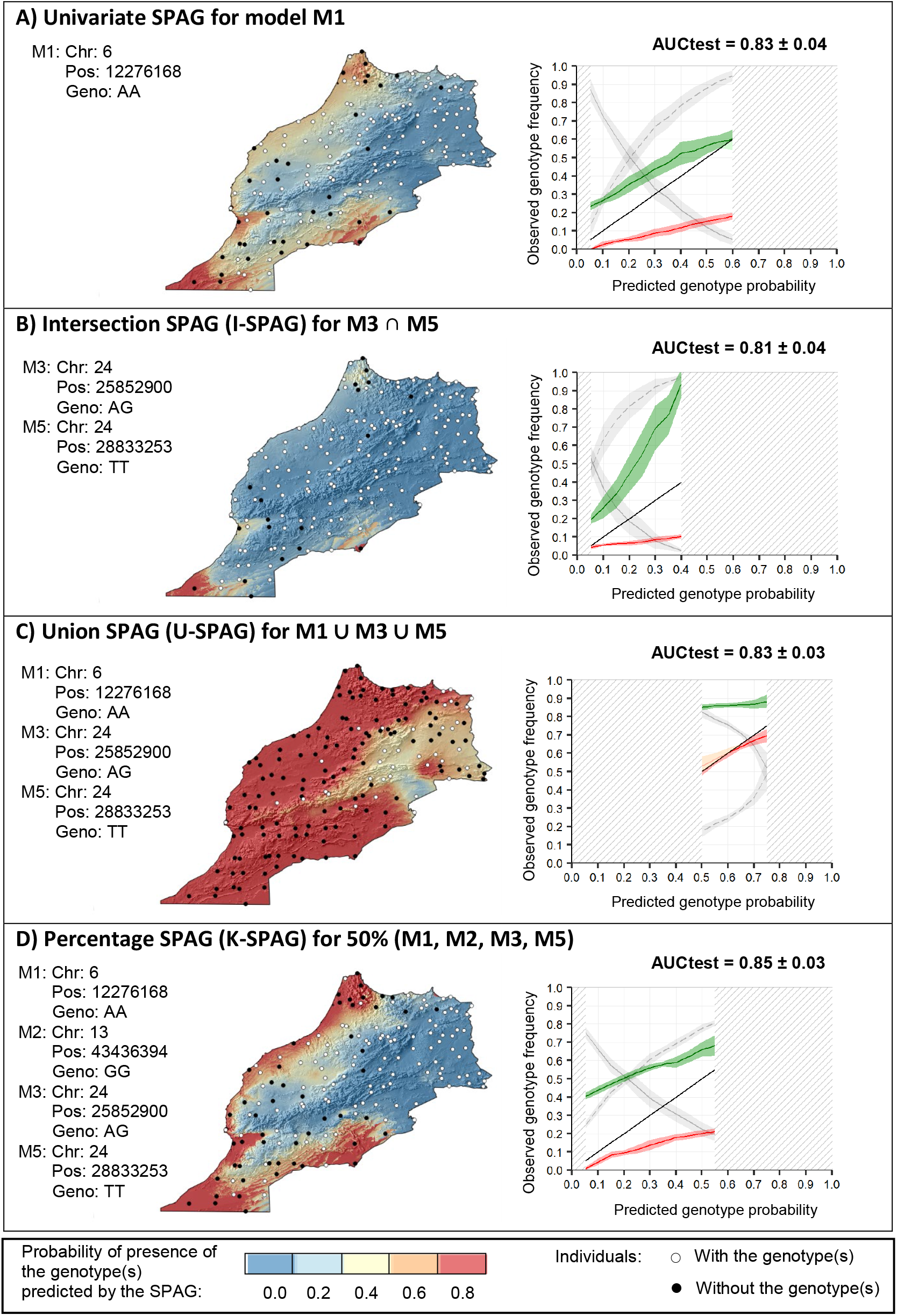
SPAG – Moroccan dataset. Univariate and Multivariate Spatial Areas of Genotypes Probability for the Moroccan dataset. The identifiers of the presented models (M1, M2, M3, M5) refers to Table 2. The maps show the average genotype(s) frequency(ies) based on the 10 runs computed with different random selection of training sets containing 25% of the total number of individuals. AUCtest indicates the mean value of the AUC computed with the testing dataset over the 10 runs. Please refer to Box 1 to interpret the validation graphs shown on the right of each map.

#### Projections under climate change

Figure 3 shows the differences between the current SPAGs presented in Figure 2 and their corresponding projections for 2070. In Morocco, the precipitation seasonality (bio15) is predicted to increase in the northwest of the country, with a maximum increase of 5 to 10% in the extreme northwestern region (Tangier-Tetouan, see region’s map in Supp. File 3) and to decrease in other areas, especially in the Atlas Mountains and near the Sahara (from −10 to −20%). The evolution of the univariate SPAG for model M1 (Figure 3A) consequently indicates the highest risk in the Tangier-Tetouan area, where the mismatch between the current and future SPAGs indicates that the probability of finding the genotype of interest should be 20% higher to find individuals well adapted to future conditions. However, many individuals in this area already carry the favourable genotype, and the risk for the population may thus be reduced thanks to natural gene flow. Nevertheless, this is not the case in the southwest of this area (Rabat, Casablanca) where the probability of finding the genotype should also be 10-20% higher according to the SPAGs difference, and none of the goats sampled there currently carry the adaptive variant. Similar observations can be made as regards the I-SPAG of M3 and M5 (Figure 3B), two other markers that may potentially confer an adaptation to high variations of precipitation. However, the U-SPAG (Figure 3C) highlights no vulnerable areas, which indicates that if the presence of at least one of the adaptive variants is sufficient to enable the adaptation to high variations of precipitation, no population may be at risk. Finally the K-SPAG (Figure 3D) also shows a risk area in the northwest of the country, where the probability of carrying the adaptive variants should be approximately 20% higher. Again, individuals in the northernmost part of this risk area may be less threatened due to the close presence of goats already carrying the favourable genotypes, whereas the population from the Rabat-Casablanca area may be more threatened due to the current much lower presence of the adapted variants.

**Figure 3.**
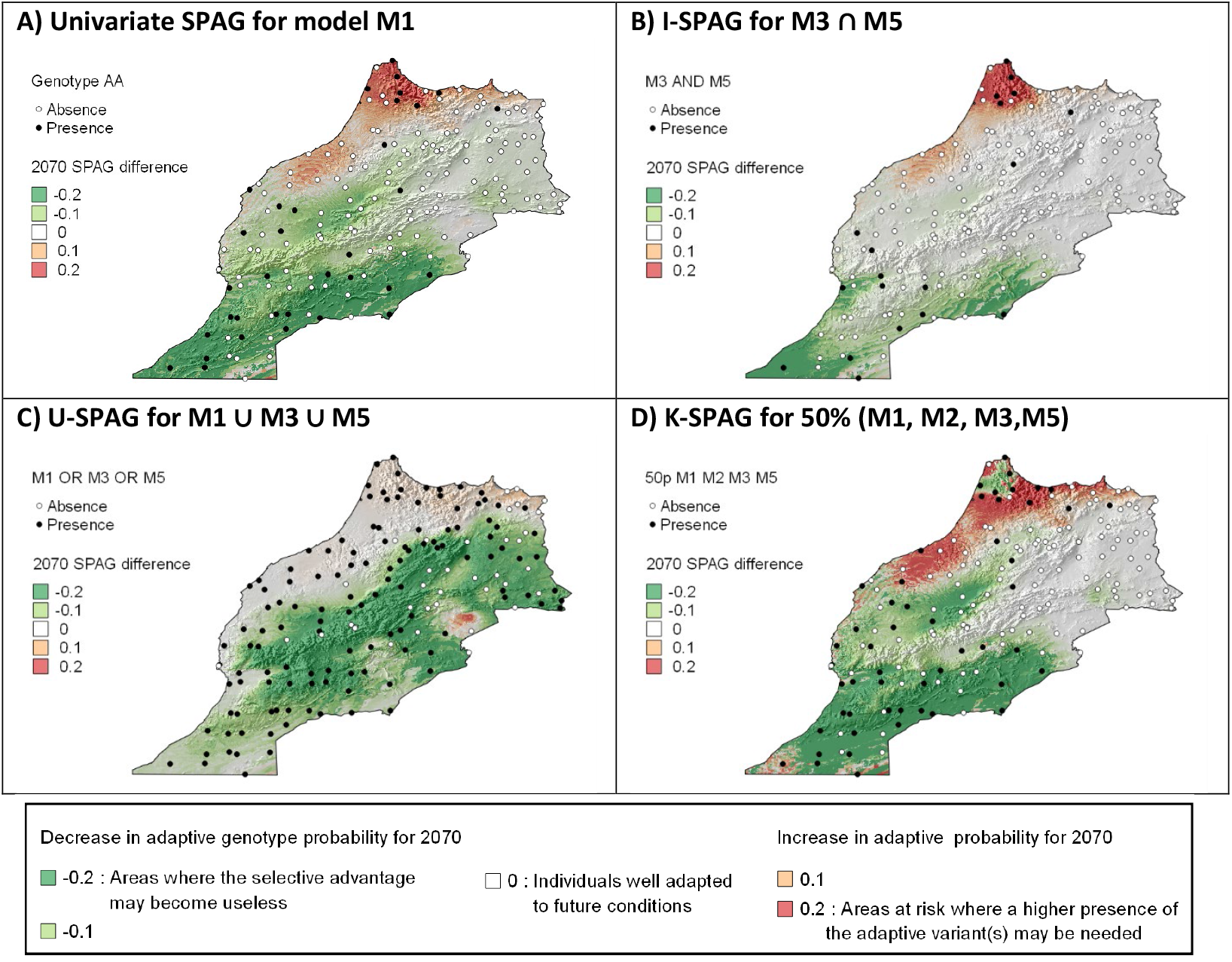
Moroccan goats - Predicted change in genotype probability for 2070. Predicted SPAG difference for 2070 considering the MPI-ESM-LR climate change scenario with RCP 8.5, for the Moroccan goats. The identifiers of the presented models (M1, M2, M3, M5) refer to Table 2. The maps show the average difference in probabilities of finding the genotype(s) based on the 10 runs computed with different selection of training sets.

### European goats

#### Population structure

For the European goats, the first component of the PCA explains 6.2% of the total variance while the second, third and fourth components explain 2.0%, 1.7% and 1.6% respectively. The low variance explained by each PCA component indicates the absence of a clear population structure. However, since the variance explained by the first component is much higher than that explained by the next ones, it is possible that the first component is partially related to the population structure. We therefore computed logistic regressions without covariates and then logistic regressions with a covariate corresponding to the coordinates of goat individuals on the first component of the PCA.

#### Logistic regressions

More than 2.6 million logistic association models were computed, of which 4.9% were significant both without covariate and with the first PCA-component as covariate, according to both G score and Wald score corrected for a false positive rate of 5% (Supp. File 6). The ten genomic regions associated with the strongest G scores when computed without covariate are presented in Table 3. The corresponding models involved two bioclimatic variables related to precipitation (bio13 = precipitation of the wettest month, bio18 = precipitation of the warmest quarter) and two bioclimatic variables related to temperature (bio3 = isothermality, bio8 = mean temperature of the wettest quarter). Seven annotated genes correspond exactly to one of the SNPs identified: KRT12, CSN1S2, CACNB2, PRDM5, LOC102174324, PALM and NAV3.

**Table 3.** Significant models – European datasets. Models corresponding to the 10 most significant genomic regions (based on G score of the model without covariate) obtained for the analysis of the European dataset, considering all bioclimatic variables. Chr=Chromosome, BP=Position in base pairs, GENO=Genotype of interest, GF=Genotype Frequency, qGpop (resp. qWpop) = FDR-corrected p-values of Gscore (resp. Wald score) of the model with the first PCA-component as covariate, qG0 (resp. qW0) = FDR-corrected p-values of Gscore (resp. Wald score) of the models without any covariate, β0 and β1 = parameters of the logistic regression without covariate, Genes = Annotated genes on the genomic region.

| ID | ENV | CHR | BP | GENO | GF | qGpop | qWpop | qG0 | qW0 | $\beta_0$ | $\beta_1$ | Genes |
| --- | --- | --- | --- | --- | --- | --- | --- | --- | --- | --- | --- | --- |
| E1a | bio18 | 19 | 40696776 | GG | 33.8 | 3.3E-09 | 3.9E-07 | 3.8E-17 | 1.2E-11 | -1.41 | 1.30 | KRT12 |
| E1b | bio13 | 19 | 40696776 | AA | 40.3 | 4.3E-09 | 5.4E-07 | 1.4E-13 | 6.9E-10 | -0.52 | -1.07 | KRT12 |
| E2 | bio18 | 1 | 38183832 | AA | 31.7 | 3.0E-09 | 1.3E-07 | 1.6E-15 | 1.2E-11 | -0.97 | 1.13 | - |
| E3a | bio3 | 6 | 86081075 | CC | 31.2 | 3.5E-11 | 1.2E-08 | 3.1E-14 | 4.2E-11 | 0.65 | -1.06 | CSN1S2 |
| E3b | bio18 | 6 | 86081075 | CC | 31.2 | 9.6E-11 | 6.9E-08 | 5.7E-14 | 5.0E-10 | 0.71 | 1.12 | CSN1S2 |
| E4 | bio8 | 13 | 32300758 | GG | 31.2 | 6.1E-10 | 9.0E-08 | 5.1E-14 | 5.6E-11 | 0.11 | -1.02 | CACNB2 |
| E5 | bio18 | 5 | 23213822 | GG | 39.0 | 6.0E-10 | 4.1E-08 | 7.3E-14 | 5.6E-11 | -0.52 | 1.02 | - |
| E6 | bio8 | 6 | 4945809 | AA | 19.4 | 9.3E-07 | 7.9E-05 | 1.0E-13 | 9.2E-08 | -2.06 | 1.55 | PRDM5 |
| E7 | bio18 | 16 | 76397454 | GG | 22.8 | 1.1E-10 | 5.3E-07 | 1.0E-13 | 8.0E-09 | 1.30 | 1.29 | LOC102174324 |
| E8 | bio18 | 7 | 67159272 | CC | 35.1 | 6.9E-10 | 1.3E-07 | 1.0E-13 | 2.3E-10 | 0.27 | 1.03 | PALM |
| E9 | bio3 | 5 | 7093719 | GG | 39.8 | 7.0E-11 | 1.8E-08 | 1.0E-13 | 9.0E-11 | -0.63 | 1.02 | NAV3 |
| E10 | bio18 | 14 | 85434737 | AA | 47.4 | 2.6E-09 | 1.6E-07 | 1.3E-13 | 2.2E-10 | -0.16 | -1.01 | - |

#### SPatial Areas of Genotype Probability

Figure4A shows the univariate SPAG for the genotype of model E10 presented in Table 3 (see Supp. File 7 for the other univariate SPAGs). This genotype is negatively associated with precipitation of the warmest quarter (bio18). The predicted probability of finding this genotype is the lowest in the Swiss Alps (<0.1), slightly higher in the Jura, French Alps and Swiss Plateau (between 0.2 and 0.4) and above 0.5 everywhere else, with a maximum around 0.8 on the Mediterranean coast. The I-SPAG (Figure 5B) shows the probability of simultaneously finding this genotype (model E10) and the genotype from model E1b, associated with low precipitation during the wettest month (bio13). The two models E1b and E10 may therefore indicate an adaptation to low values of precipitation. However, the I-SPAG indicates that their simultaneous presence is not very likely (predicted frequency < 0.6 everywhere). The predicted probability is the highest in the centre-north of France (regions Centre, Iles de France, East of Pays de la Loire, Normandie and South of Hauts de France, see Supp. File 3 for regions’ map) and in the southern part (Occitanie and West of Provence), whereas in the Alps, Jura and most of Switzerland, the probability of simultaneously finding these two genotypes is close to 0 and none of the sampled goats carry them both. The probability of finding at least one of these two variants, presented on the U-SPAG in Figure 4C, indicates a trend similar to the probability of presence of E10 alone (Figure 4A), but with even stronger contrast between the Alps-Jura-Switzerland area (frequencies < 0.3) and the rest of the territory (frequencies > 0.7). Finally the K-SPAG (Figure 4D) shows the probability of finding at least 50% of five genotypes negatively associated with precipitation in the warmest quarter (bio18), i.e. the probability of finding at least three of them. Note that for the models positively associated with bio18 in Table 3, we used the alternative genotype that was the most significantly negatively correlated with bio18 (indicated after the model ID). The resulting SPAG is very close to the I-SPAG of E1b and E10 (Figure 4B). For all cases presented, the validation graphs indicate that the SPAGs computed with 25% of the individuals generally correctly predict the genotype frequency of the remaining 75% of individuals.

**Figure 4.**
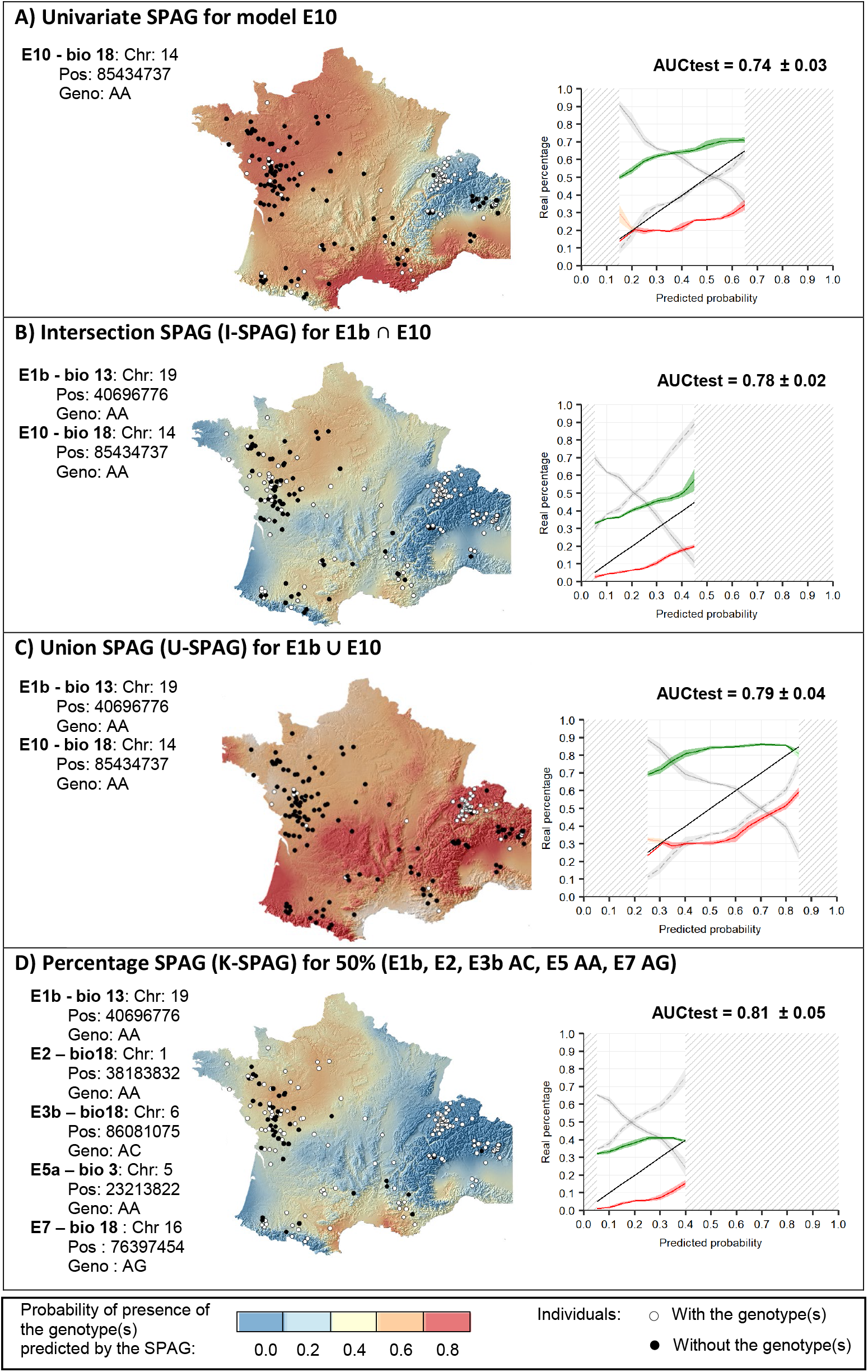
SPAG – European dataset. Univariate and Multivariate Spatial Areas of Genotypes Probability for the European dataset. The identifiers of the presented models (E10, E1d, E2, E3b, E5, E7) refers to Table 3. The maps show the average genotype(s) frequency(ies) based on the 10 runs computed with different random selection of training sets containing 25% of the total number of individuals. Note that since up to five individuals can be localised on the same site, a black dot indicates a presence if at least 50% of the individuals of the site carry the marker(s). Please refer to Box 1 to interpret the validation graphs shown on the right of each map.

**Figure 5.**
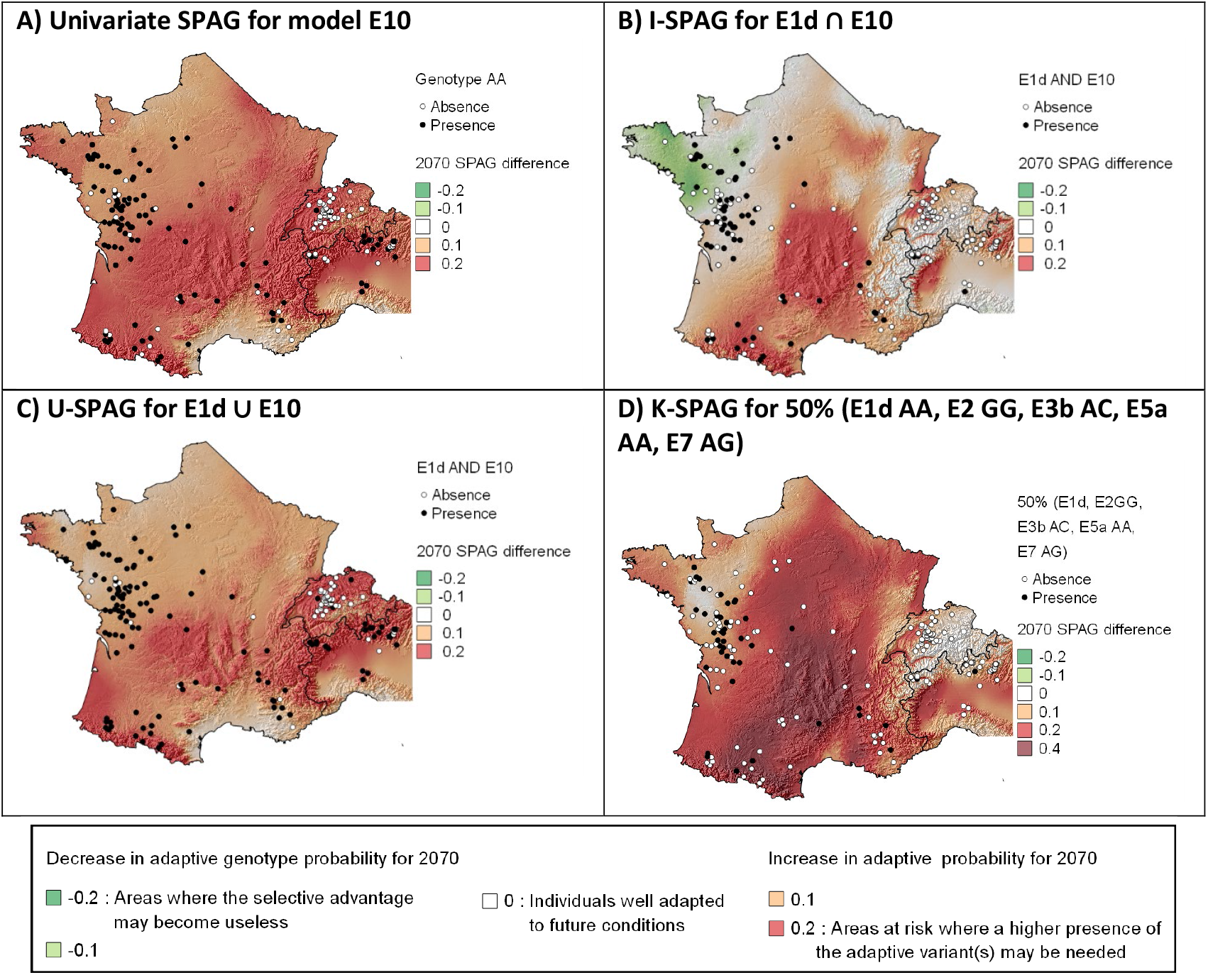
Predicted change in genotype probability for 2070. Predicted change in genotype probability for 2070 considering the MPI-ESM-LR climate change scenario with RCP 8.5, for the European goats. The identifiers of the presented models (E10, E1b, E2, E3b, E5a, E7) refer to Table 3. The maps show the average difference in probability of finding the genotype(s) based on the 10 runs computed with different random selection of training sets.

#### Projection under climate change

Figure 6 shows the differences between the current SPAGs presented in Figure 5 and their corresponding projections for 2070. The model E10 presented in Figure 5A was related to the precipitation of the warmest quarter (bio 18), which is projected to decrease by 20 to 180 mm over the whole study area until 2070. The projected precipitation loss is maximum in the Alps (−120 to −180 mm), on the Mediterranean coast (−100 to −130) and in the centre-south of France (West of Auvergne and East of Nouvelle Acquitaine, −90 to −120 mm, see region’s map in Supp. File 3). As a consequence, the mismatch between current and future SPAGs for model E10 (Figure 5A) indicates that all populations may be threatened by climate change, since the probability of finding the adaptive genotype should be 10 to 20% higher everywhere. None of the goats living in the Alps and Switzerland currently carry this adaptive genotype and these populations may thus be particularly vulnerable. When considering the I-SPAG of E10 and E1b (Figure 5B), a higher risk is highlighted in the centre-south of France (West of Auvergne and its surrounding), northeast of France (Alsace in East of Grand-Est), Swiss Plateau, and northwest of Italy. In centre-south of France and Swiss-Plateau, almost none of the goats sampled carry simultaneously the two genotypes and the populations may therefore be particularly threatened. The U-SPAG of the same genotypes (Figure 5C) shows results very similar to the univariate SPAG for E10 alone (Figure 5A). If the presence of at least one of these two genotypes may be sufficient to allow adaptation to low precipitation, then the most threatened regions will again be the Alps and Switzerland, where both markers are currently absent. Finally, the evolution of the K-SPAG (Figure 4D) shows a high risk in a large part of France and northern Italy, where the SPAG’s mismatch indicate that the probability of finding the adaptive genotypes should be more than 20% higher.

## Discussion

### Mapping genotype probabilities

The first utility of SPAGs is to quantify the current probability of finding beneficial genotypes or the expression of favourable traits in plant and animal populations, even in regions where no individuals have been sampled. Our results show that with few training individuals (i.e. 50 simulated individuals, 41 Moroccan or 96 European goats), a good estimate of the probability of finding genotype(s) of interest is possible. The univariate models presented here have already been applied to map the genotype frequencies of adaptive variants of the Scandinavian brown bears (Joost, 2006), Moroccan sheep (Rochat *et al.,* 2016) and coral reefs (Selmoni *et al.*, 2019). Multivariate models are presented here for the first time and, according to the validation procedure applied on a simulated dataset and two case studies, they appear to be powerful in estimating the combined probability of finding several genotypes potentially correlated with different environmental variables. With the I-SPAG, the resulting probabilities may rapidly become very low, but this model could be used when we suspect that the simultaneous presence of some adaptive genotypes is needed to ensure the adaptation, or when we would like to highlight the probability of simultaneously finding variants that may confer an adaptive response to different environmental variables (e.g. low precipitation and high temperature). At the other extreme, the U-SPAG may rapidly indicate high probabilities of presence in most parts of the territory, but it can be used when it is suspected that the presence of at least one of the genotypes may be sufficient to confer adaptive potential. Since it is generally difficult to know whether the simultaneous presence of adaptive genotypes is needed or whether an union is sufficient, K-SPAG offers an interesting compromise, allowing the identification of populations that retain a given percentage of variants, which can help delineate areas where there is the highest probability of finding individuals with high adaptive potential.

### From SPAG to conservation

The study of the shift in SPAGs under climate change conditions can help identify 1) well-adapted populations, where individuals currently show adaptive genotypes that seem to be optimal under future conditions, 2) populations at risk where the current genotype frequency is not optimal, but where the favourable genotypes are already present in the population, thus potentially allowing a natural increase in genotype frequency through gene flow and 3) threatened populations where optimal variants are currently lacking but would be needed to ensure adaptation to future climate. These identifications may be of great value for conservation planning. Indeed, when prioritising conservation areas, success could be enhanced by choosing to preserve preadapted individuals that already carry functional variants conferring them good adaptation to future climate (Orr and Unckless, 2008). Moreover SPAGs can also be used to prevent the translocation of individuals that do not currently carry the variants favourable for future conditions at the target site, which would result in a reduction or loss of adaptive potential of the target populations (Weeks *et al.*, 2011). In addition, conservation plans can be developed to increase the survival capacity of threatened populations. This may involve assisted gene flow to import adaptive variants into a population where they are lacking (Aitken and Whitlock, 2013; Kelly and Phillips, 2016) or artificial selection of individuals already pre-adapted to future conditions (Hoffmann, 2010). However, this has to be undertaken carefully since the selection of locally adapted individuals can result in a loss of genetic diversity (Savage *et al.*, 2018), which may decrease the potential of populations to adapt to new environmental changes. Kardos and Shafer (2018) therefore proposed that gene-targeted conservation measures should only be taken with traits affecting vital processes of the species and when phenotypic variation is large enough to ensure a high probability of success. Finally, the SPAG maps presented could be integrated into decision-making frameworks considering the adaptive potential when defining the vulnerability of species (Bonin *et al.*, 2007; Williams *et al.*, 2008; Sgrò *et al*., 2011; Dawson *et al.*, 2011; Razgour *et al.*, 2018) or in more global decision frameworks that take into account other vulnerability factors such as the predation level or habitat loss.

### The goat’s example

Several signatures of local adaptation where identified for the goats under study. In Morocco, three of the genes identified (DSG4, KCTD1 and CDH2) may be related to the development of hair (Kljuic *et al.*, 2003; Ling *et al.*, 2014; Wang *et al.*, 2017; Zhang *et al.*, 2019) or skin properties (Hayashi *et al.*, 2007). These results suggest that goats confronted with high variations of precipitation may have adapted through a modification of hair or skin traits, which could for example ensure a better water repulsion. In Europe, two of the genes highlighted as potentially conferring an adaptation to drought conditions (KRT12 and PRDM5) may be related to properties of the cornea (Kao *et al.*, 1996; Burkitt Wright *et al.*, 2011). They could thus potentially highlight an adaptation to higher UV-radiation associated with driest conditions. The other genes identified are related the casein content of the milk (CSN1S2, Ramunno *et al*., 2001), the calcium channel and energy pathway (CACNB2, Cardona *et al.*, 2014) or the skin properties (PALM and NAV3, Kutzleb *et al.*, 1998; Karenko *et al.*, 2005). Many of the genes highlighted on the two case studies may therefore be associated with a function that can be influenced by climate, which reinforces the potential that they are true signatures of local adaptation. However, it is known that logistic regressions such as implemented here, as most of the other methods, may lead to the identification of false positive (Stucki *et al.*, 2017), and the results should thus be confronted with other methods available to detect signatures of natural selection. In addition, although previous studies show the power of genotype-environment associations to predict phenotype (Lasky *et al.*, 2015; Vangestel *et al.*, 2018) or fitness (Fournier-Level *et al.*, 2011; Hancock *et al.*, 2011), more investigations are needed to verify that the genotypes identified are really conferring an adaptive advantage (Funk *et al.*, 2019).

The Moroccan case study highlighted that goat populations from the surroundings of Rabat and Casablanca may lack adaptive genotypes potentially conferring an advantage to face high variation of precipitation. If the adaptive role of these genotypes is validated, goat populations in this region may be threatened. Because of the great economic and social importance of goats in Morocco, it is crucial to preserve viable populations. Indeed, in this country, agriculture contributes to 12 to 24% of the national GDP and employs 40% of the total active population (Boujenane, 2005). Livestock farming, especially small ruminants, is the most important sector of agriculture and goat farms account for 20% of the total number of farms (Boujenane, 2005). It is therefore important to consider preserving or introducing the adaptive genotypes on each vulnerable population. This could be done for example by favouring crossbreeding with individuals from the southern or north-eastern part of the country, where adaptive genotypes are currently well present, and by avoiding breeding or translocation with exotic goats or goats from the Atlas or Oriental regions. In the northern part of Morocco (Tanger-Tetouan regions), goat populations represent 12% of the national goat populations (Chentouf, 2014), and they play an important role in preserving food security (Godber *et al.*, 2016). In this region, crossbreeding with exotic breeds has been introduced to improve milk production (Boujenane, 2005; Godber *et al.*, 2016). However, our results show that the frequency of adaptive genotypes should increase in the goat populations from this region, and that it is therefore essential to maintain local individuals with the necessary adaptive genotypes.

### Limitations and perspectives

The SPAG approach presented appears to be powerful for mapping the probability of finding locally adapted genetic variants in a landscape. However, the adaptation process is complex and often involves polygenic traits (Pritchard and Rienzo, 2010), for which the detection power of the genotype-environment associations may be reduced (Villemereuil et al., 2014; Harrisson et al., 2014). In this case, it may be advisable to use multivariate genotype-environment association models (Forester et al., 2017) or to integrate other methods to identify SNPs related to polygenic adaptation (Zhou et al., 2013; Lasky et al., 2015). In addition, since the results of the shift under climate change may be highly dependent on the climate change scenario considered, computations should be performed with various scenarios and less weight should be given to the conclusions not consistent within scenarios (Reside *et al.*, 2018). Finally, in order to assess the real vulnerability of populations, an analysis of connectivity should be carried out to highlight the potential of natural gene flow to increase the probability of finding favourable genotypes in threatened populations.

SPAGs could also be used to predict the presence of genotype(s) associated with other pressures showing a spatial distribution, such as the presence of a parasite (Vajana *et al.*, 2018) or a predator (Cousyn *et al.*, 2001) or the urbanization level (Harris and Munshi-South, 2016). Very similar models can also be derived to predict allele frequencies instead of genotypes frequencies or to integrate other covariates (e.g. to consider autocorrelation or to use other indicators of population structure such as the Admixture coefficients). In addition, SPAGs are provided as maps, which enables a visual identification of threatened populations and could thus facilitate discussions between different conservation actors. SPAGs therefore constitute a valuable tool to support conservation decisions, especially under current changing climatic conditions.

## Code availability

The R functions developed to compute the univariate and multivariate SPAGs are available at : https://github.com/estellerochat/SPAG

