## Supplementary material for "Spatial Areas of Genotype Probability (SPAG): predicting the spatial distribution of adaptive genetic variants under future climatic conditions": Supp. File 7

*Supp. File 7 – Univariate SPAG*

Figures on the following pages presented the Univariate Spatial Areas of Genotype Probability for all models from Table 2 (Morocco) and Table 3 (Europe) presented in the main text. The maps show the average genotype(s) frequency(ies) based on the 10 runs computed with different random selection of training sets containing 25% of the total number of individuals. Please refer to Box 1 in main text to interpret the validation graphs shown on the right of each map and refer to Supp. File 2 for the list of bioclimatic variables.

Legend valid for all maps

Genotype

- Presence
- Absence

Predicted Genotype Frequency

- 0.0
- 0.2
- 0.4
- 0.6
- 0.8

### Chromosome 6: 12276168 AA

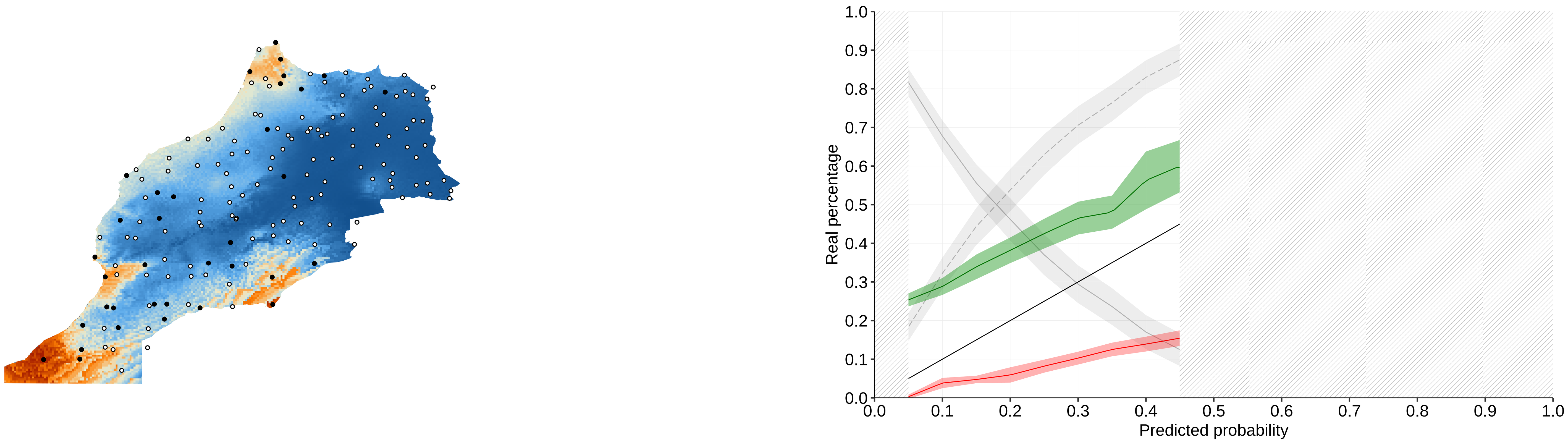


### Chromosome 13: 43436394 GG

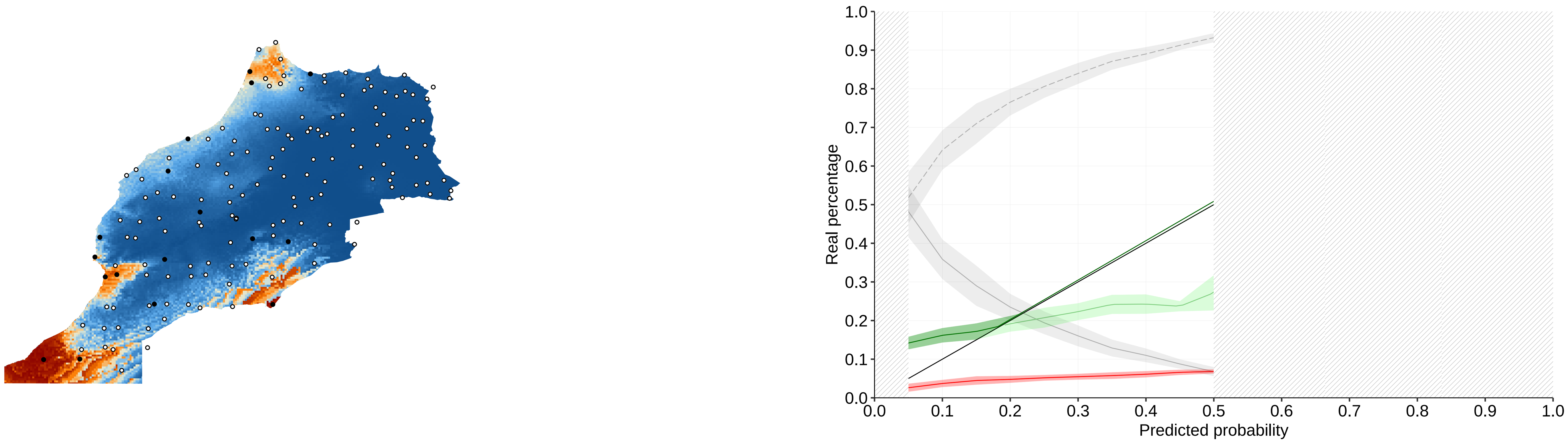


### Chromosome 24: 19436980 CC

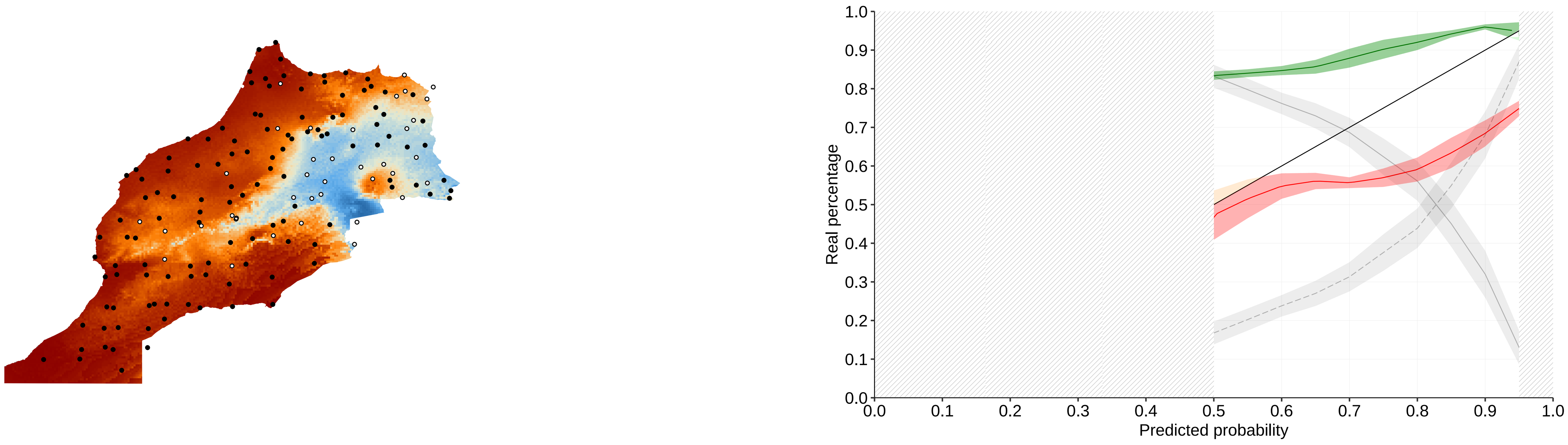


### Chromosome 24: 25860754 GA

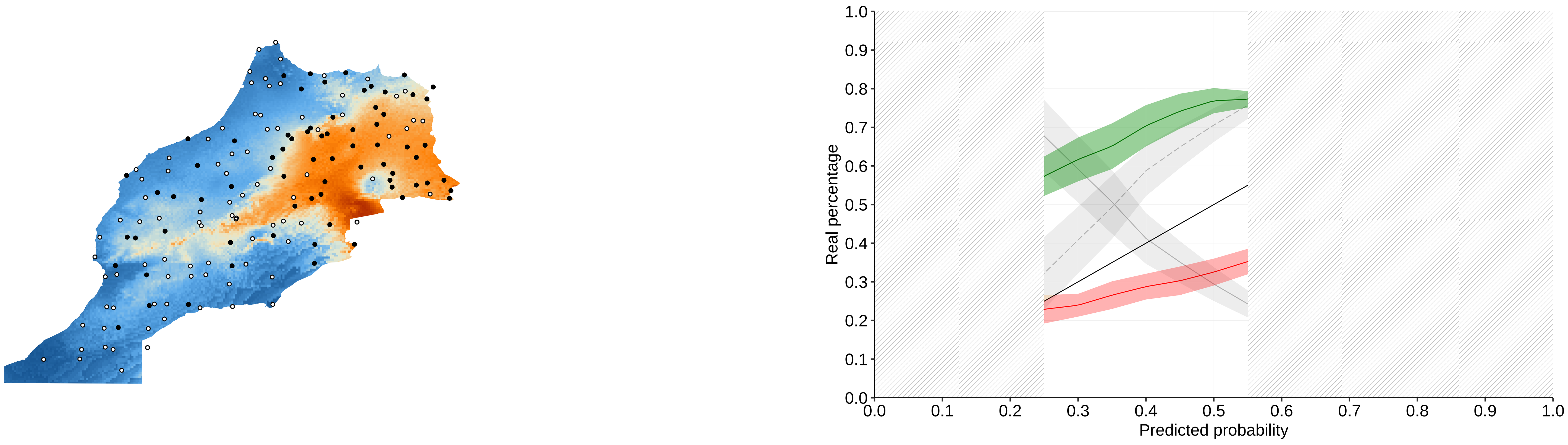


### Chromosome 24: 28833253 TT

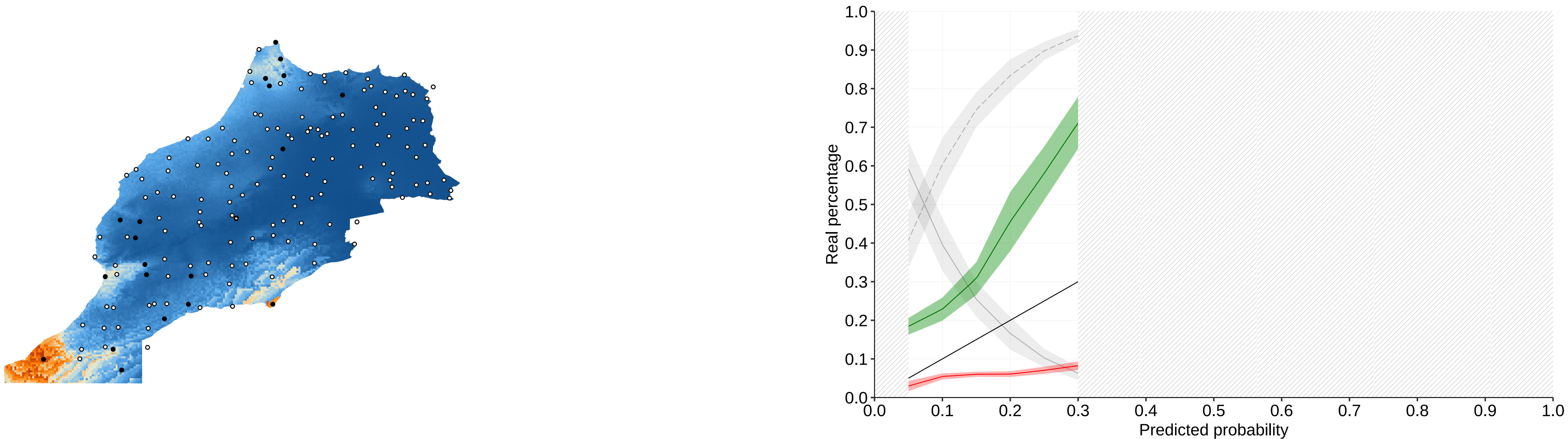

### Chromosome 24: 30566869 TT

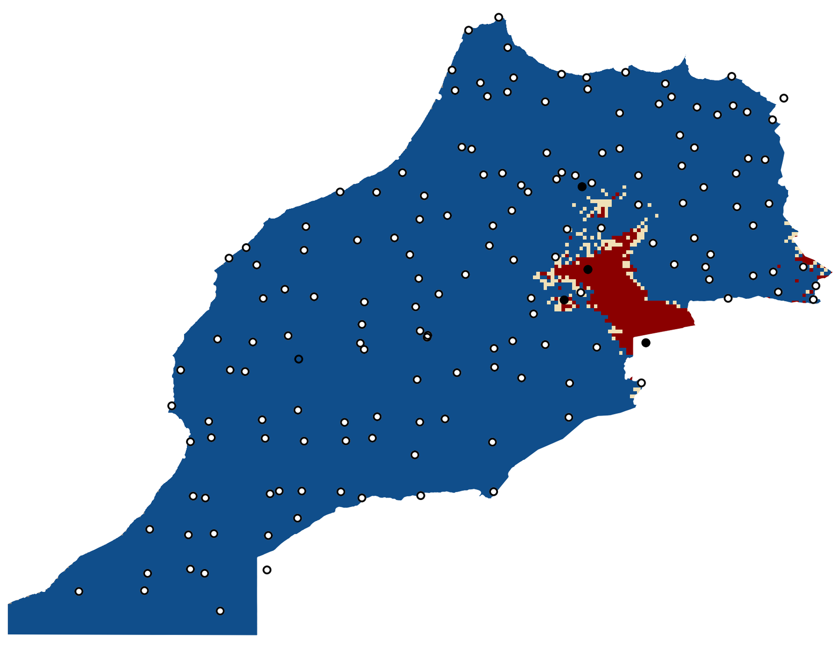

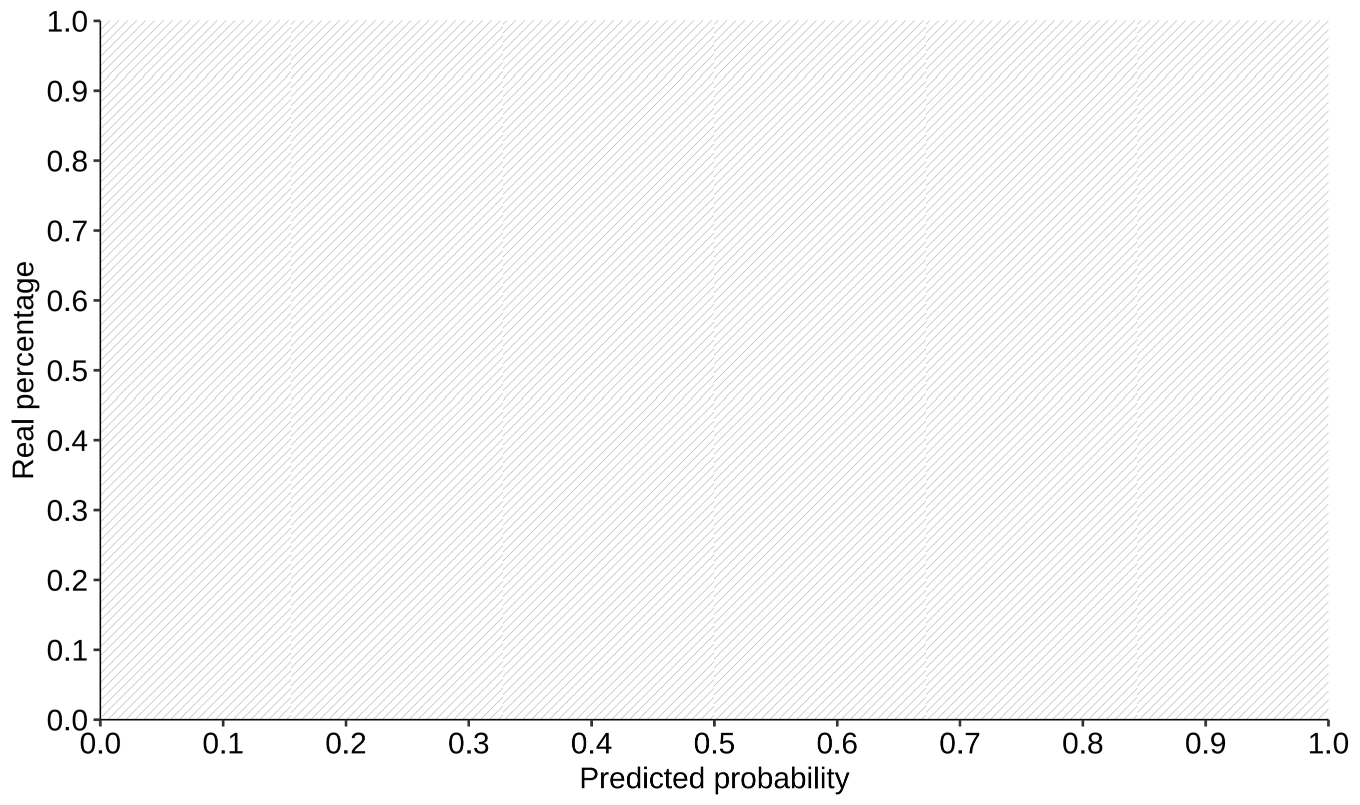


### Chromosome 27: 25930079 GG

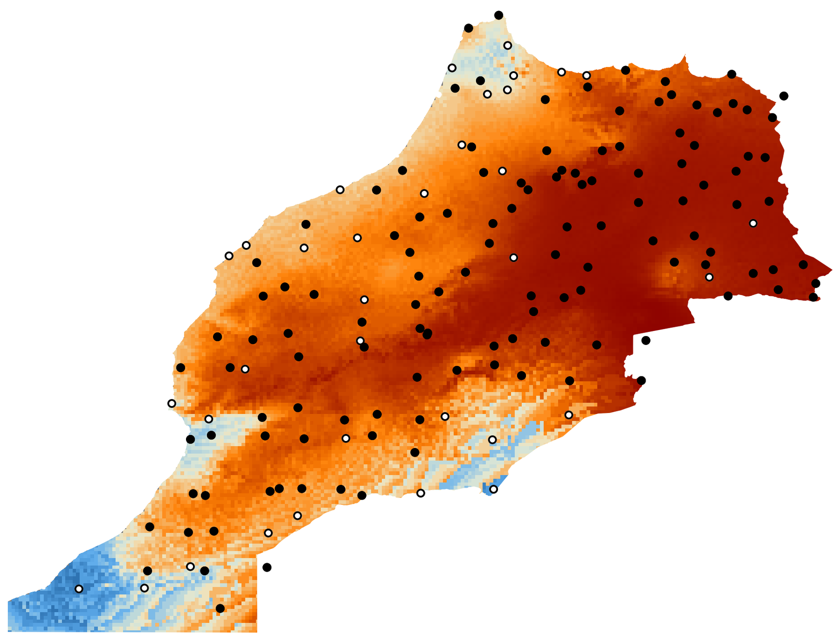

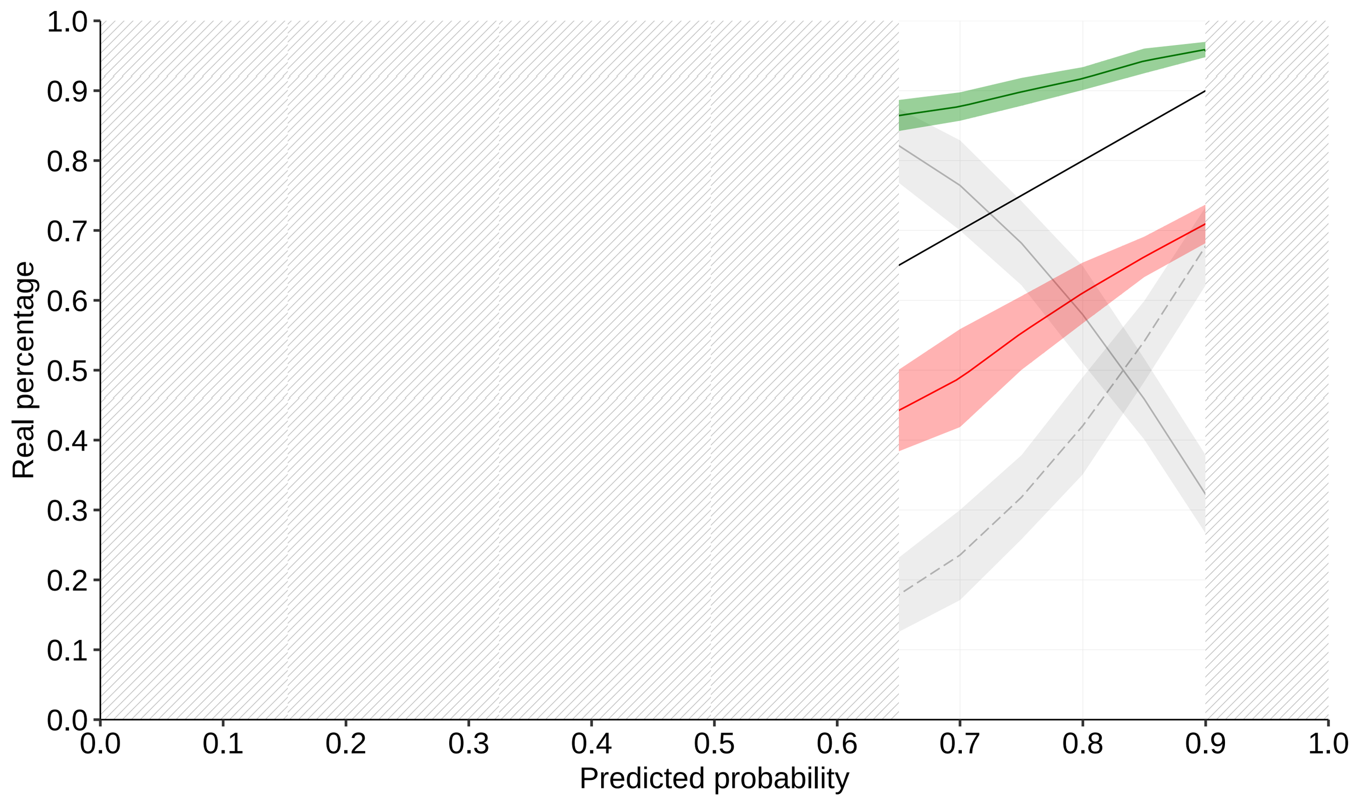

**Chromosome 19: 40224821 GG Bio 18**

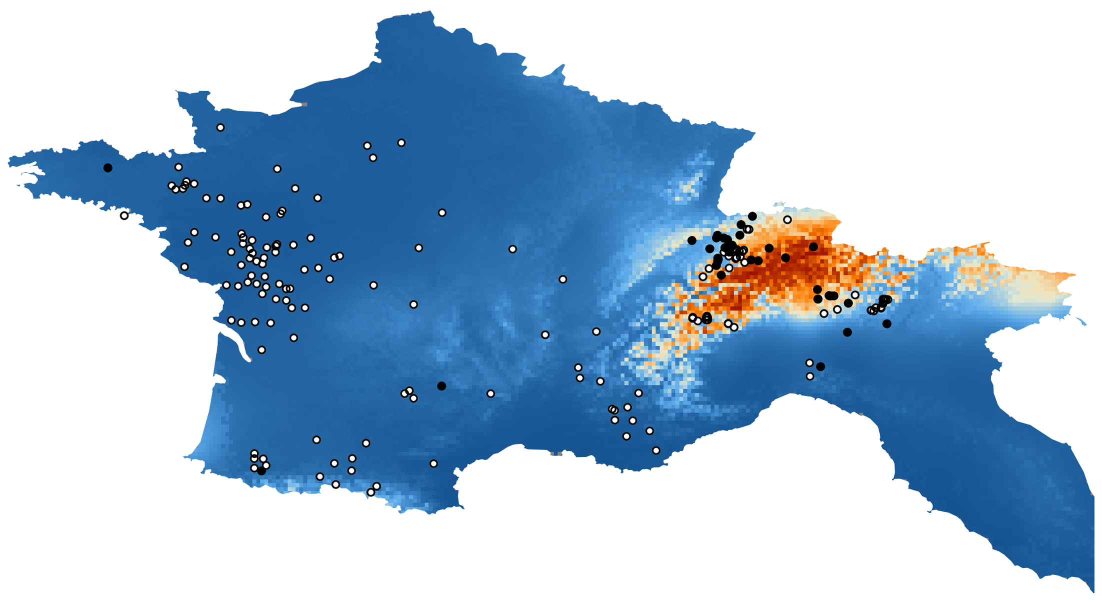

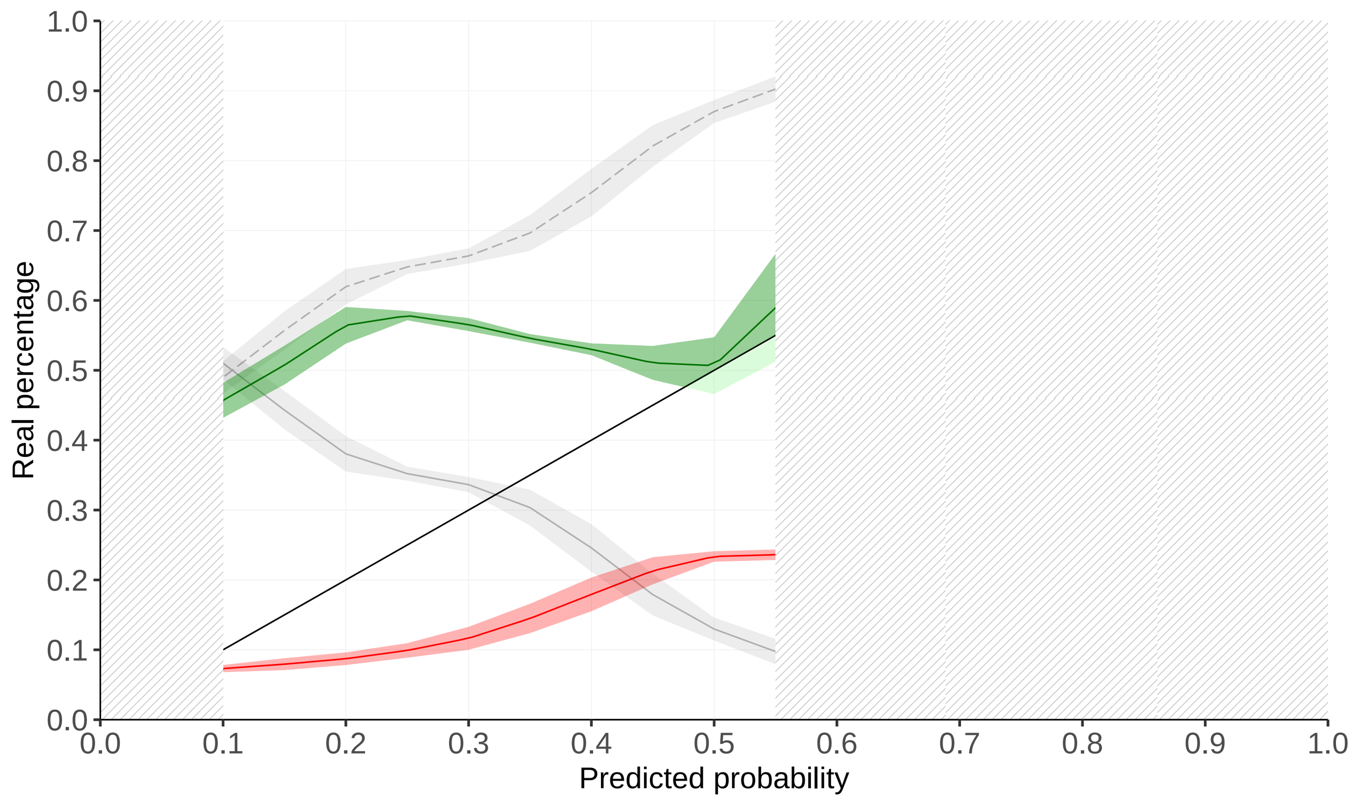


**Chromosome 19: 40224821 GG Bio 13**

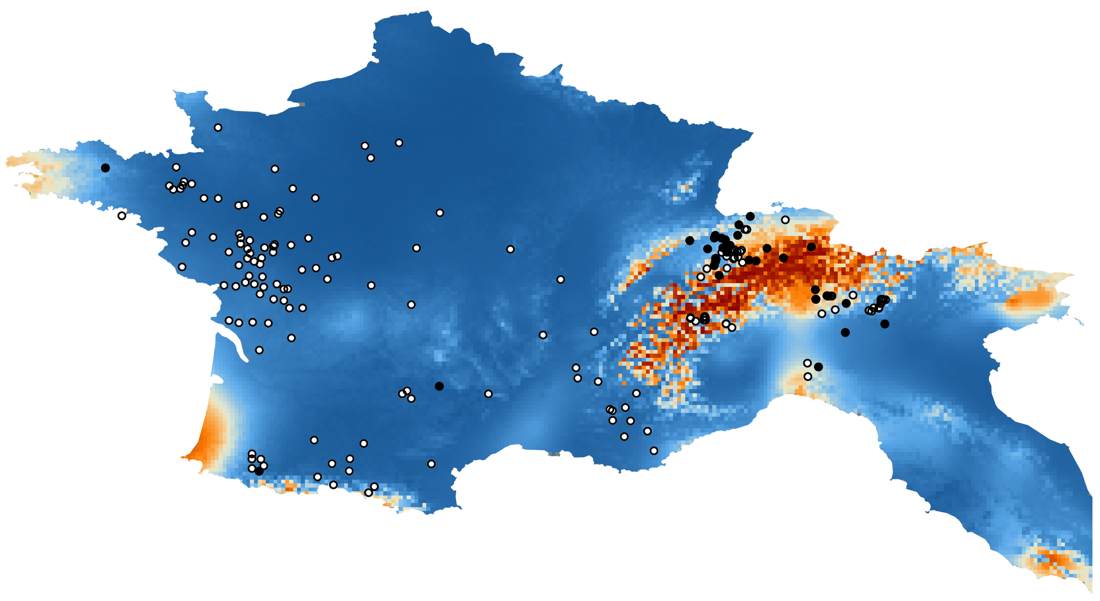

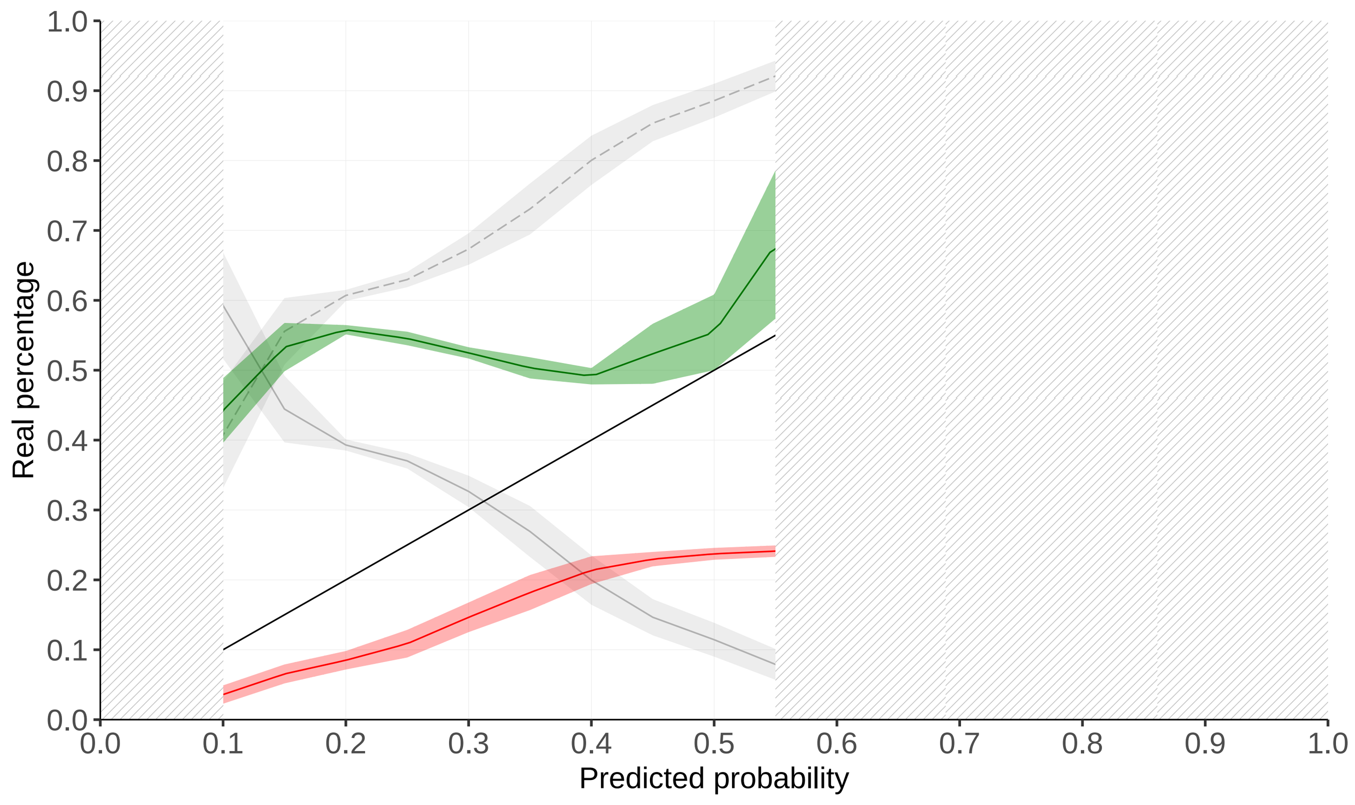


**Chromosome 19: 40224821 GG Bio 16**

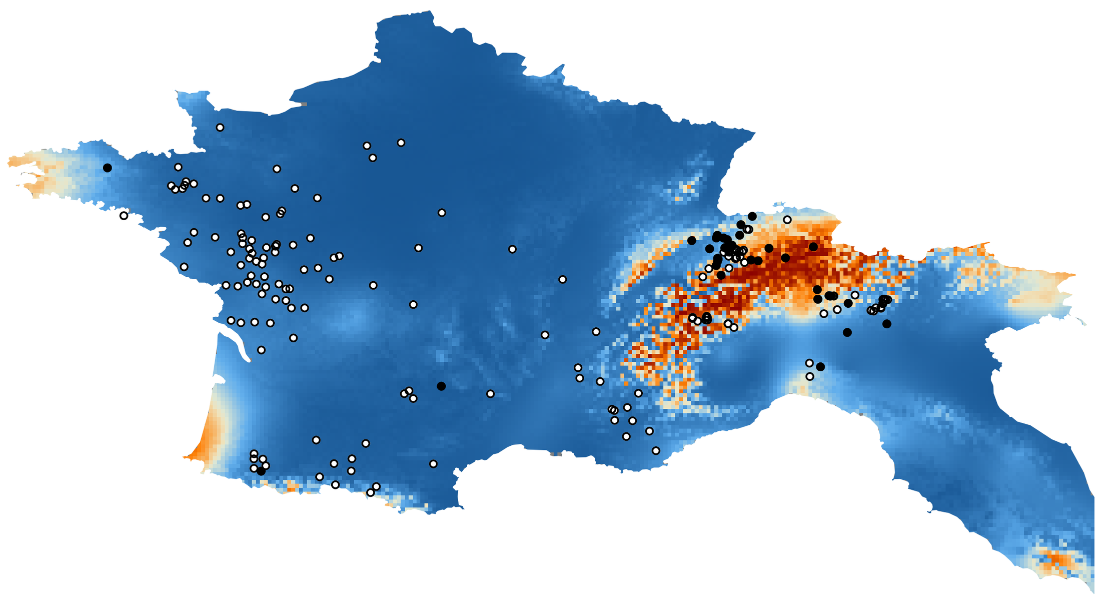

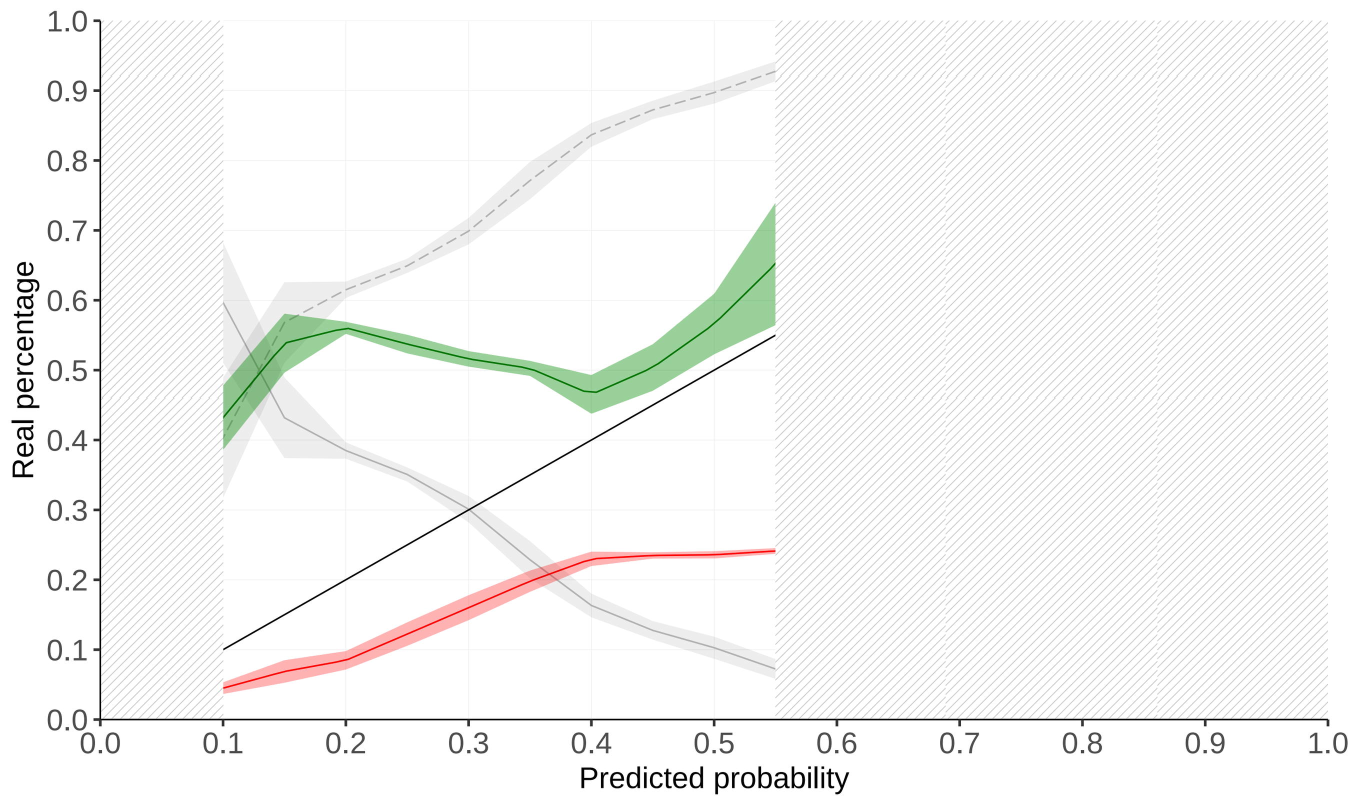


**Chromosome 19: 40224821 AA Bio 13**

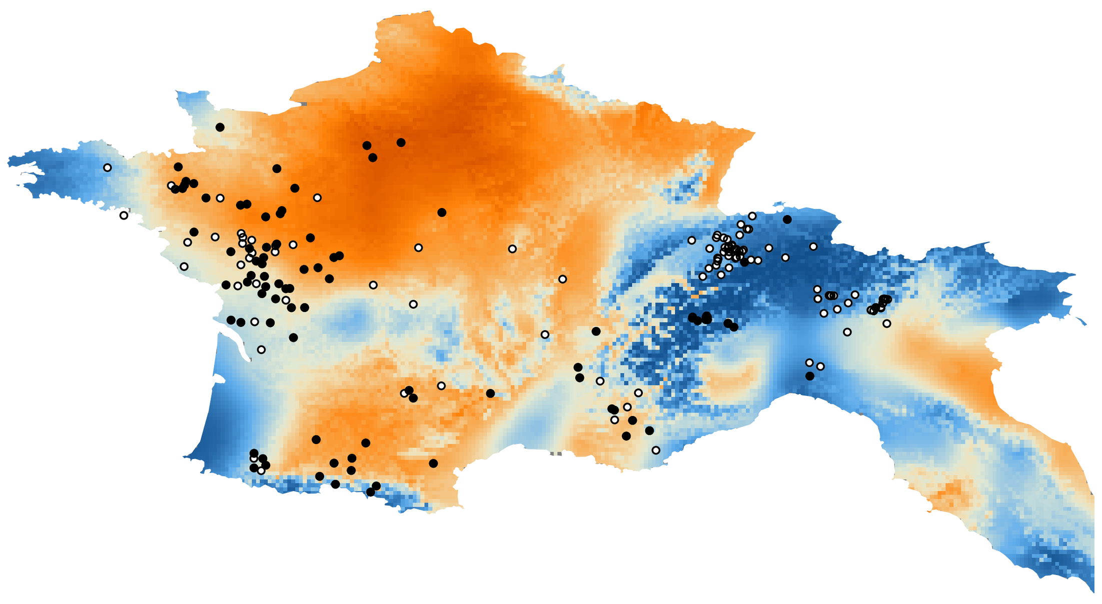

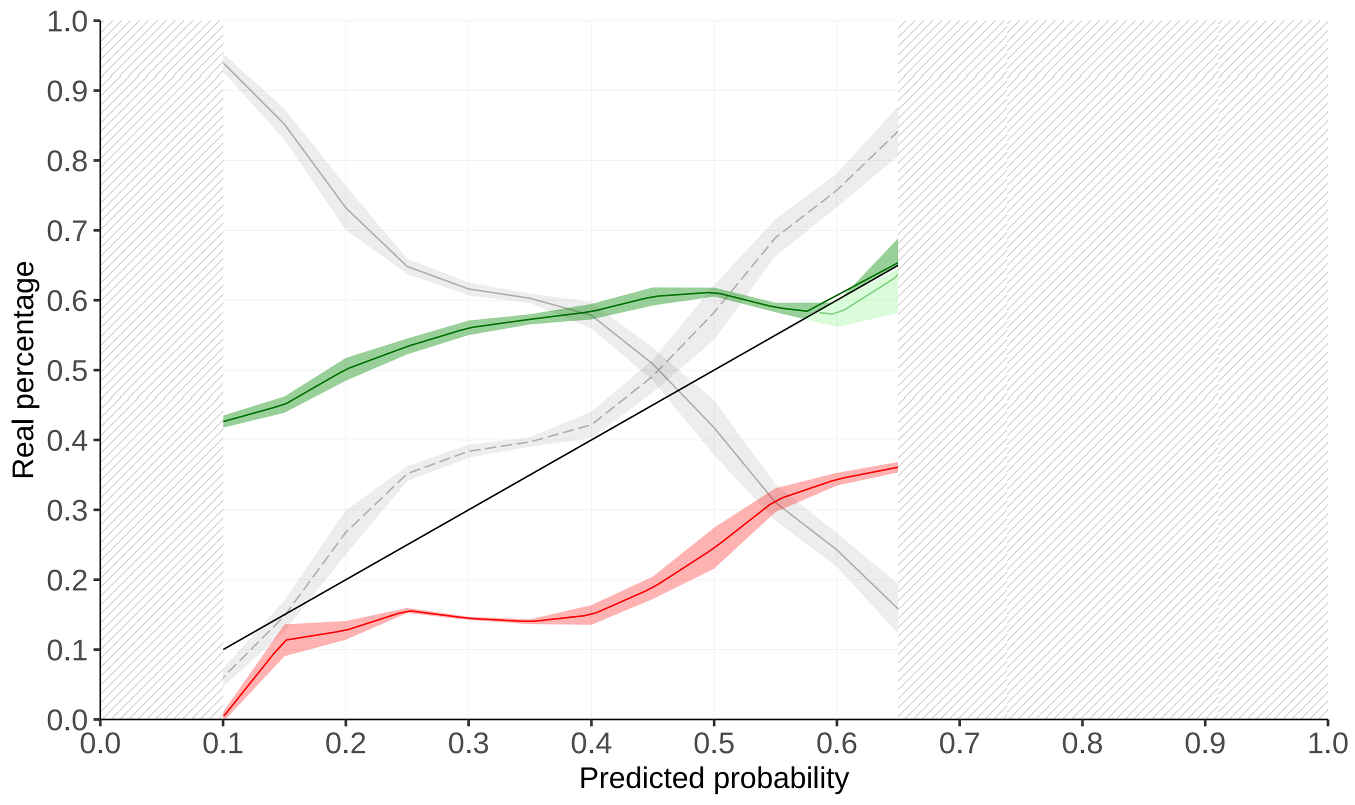


**Chromosome 1: 38282037 AA Bio 18**

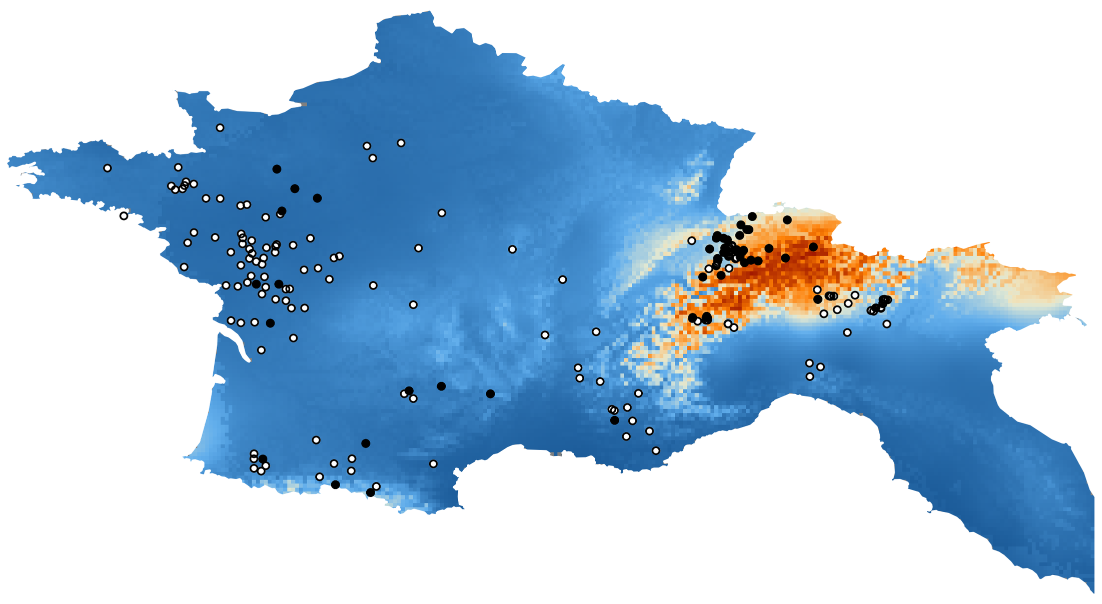

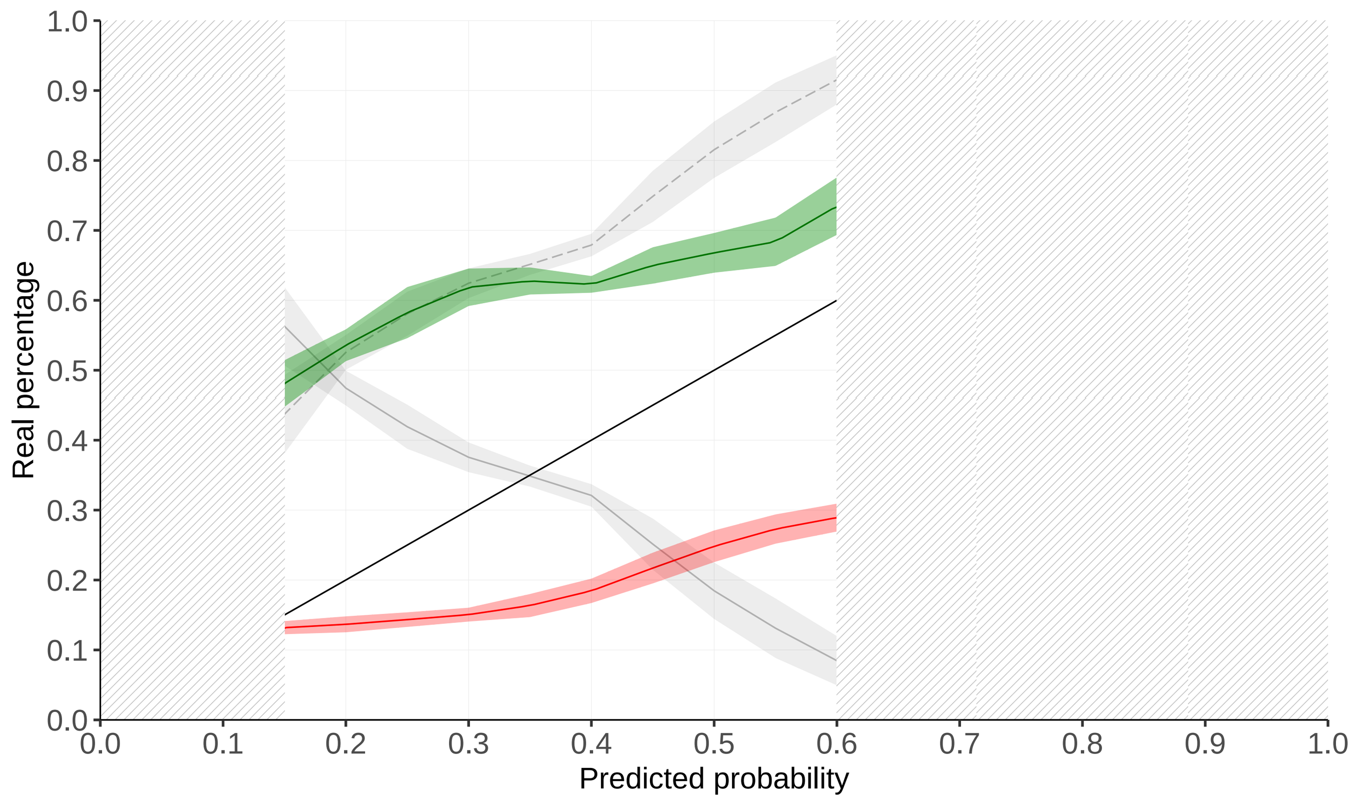

**Chromosome 6: 82779273 CC Bio 3**

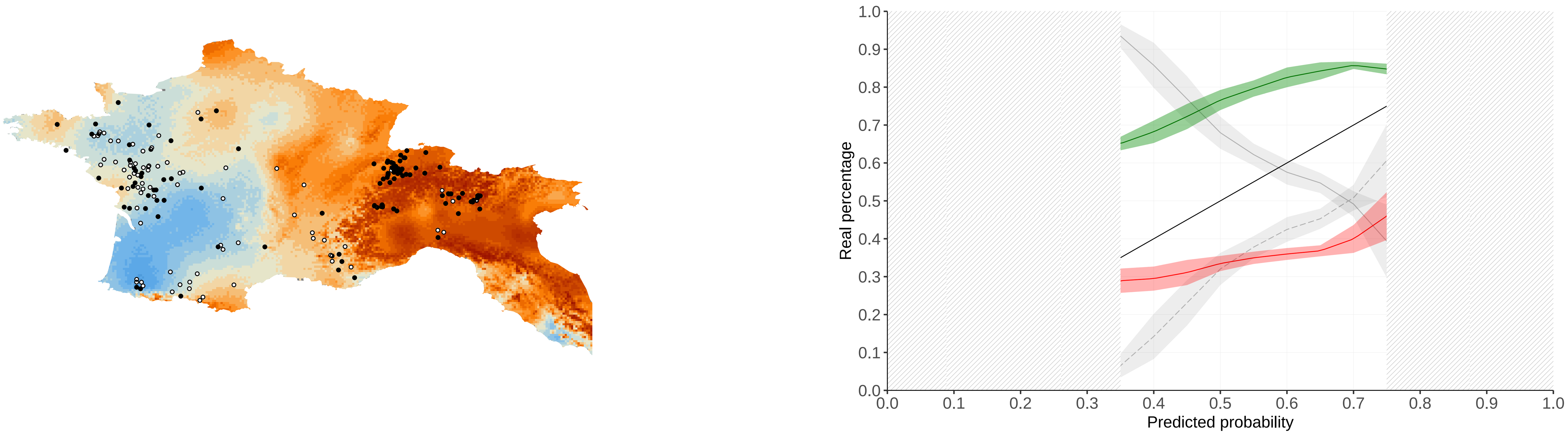


**Chromosome 6: 82779273 CC Bio 18**

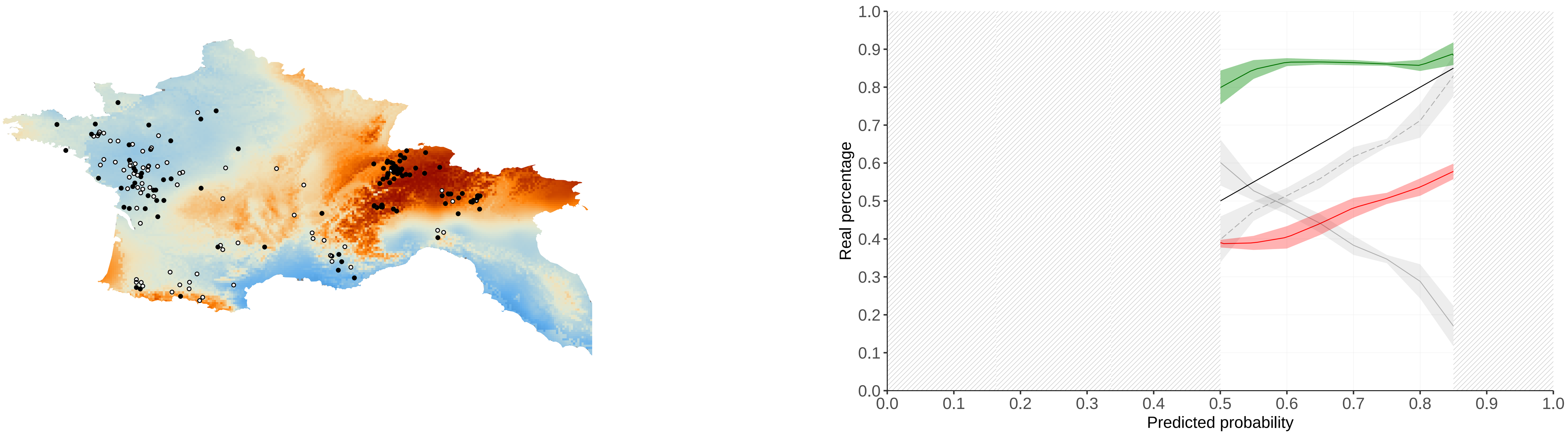


**Chromosome 13: 31676938 GG Bio 8**

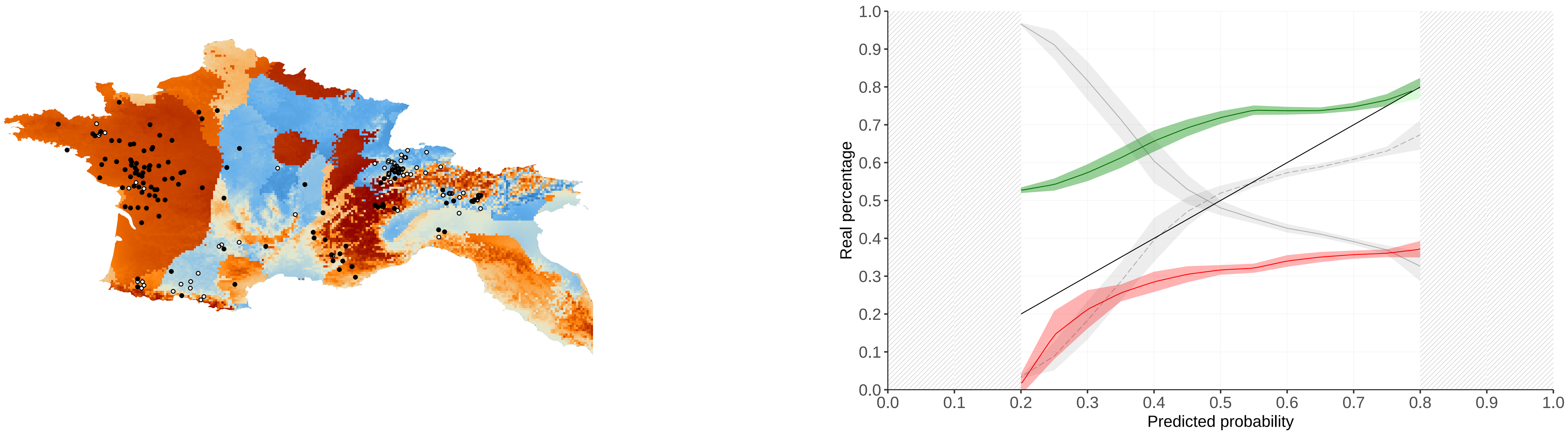


**Chromosome 5: 22749602 GG Bio 18**

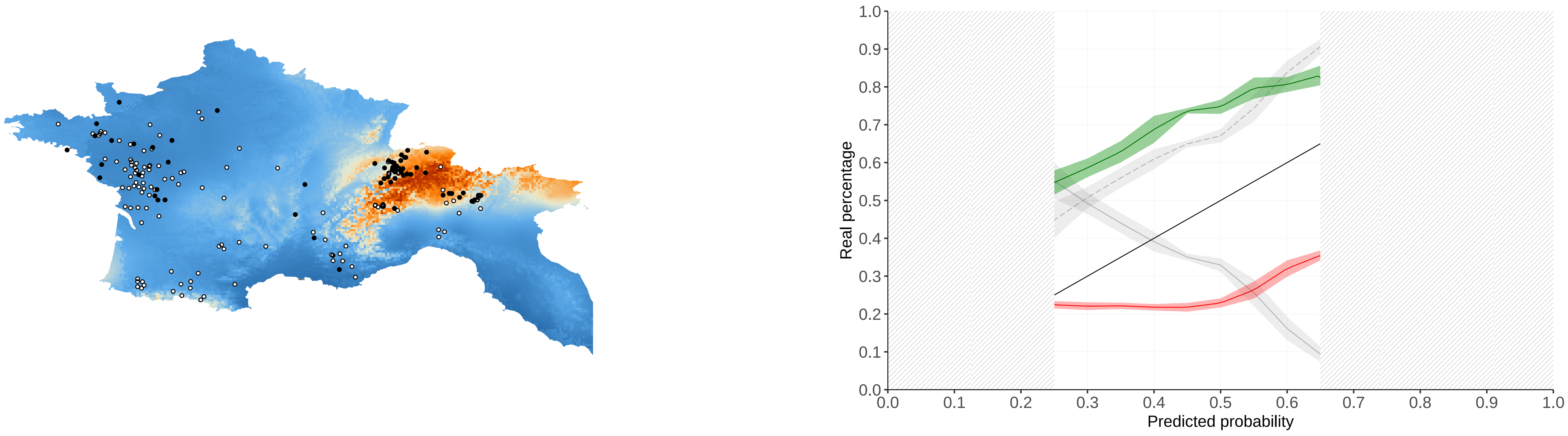


**Chromosome 5: 22749602 GG Bio 16**

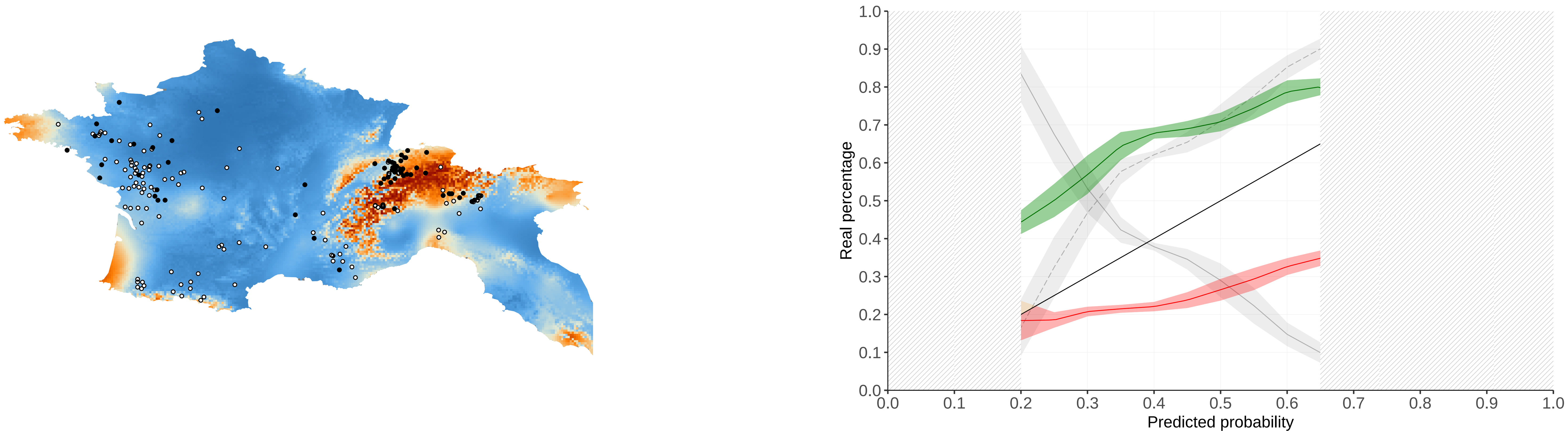

#### Chromosome 6: 4000730 AA Bio 8

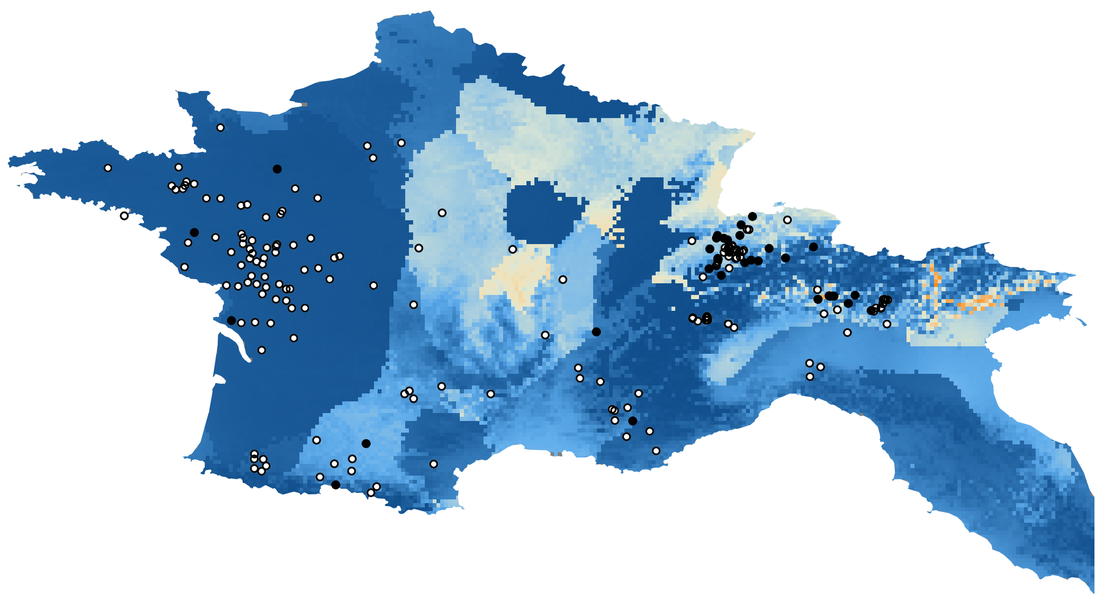

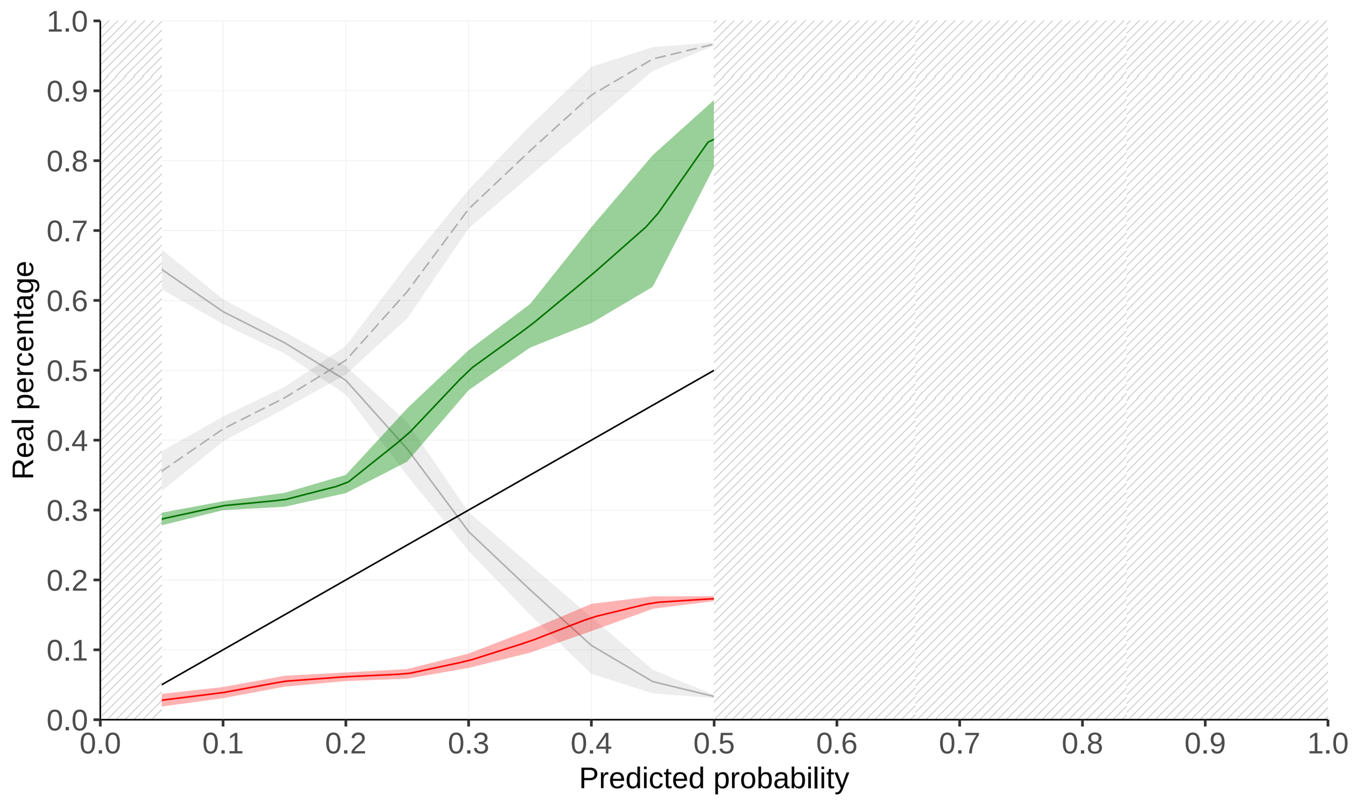


#### Chromosome 16: 75413714 GG Bio 18

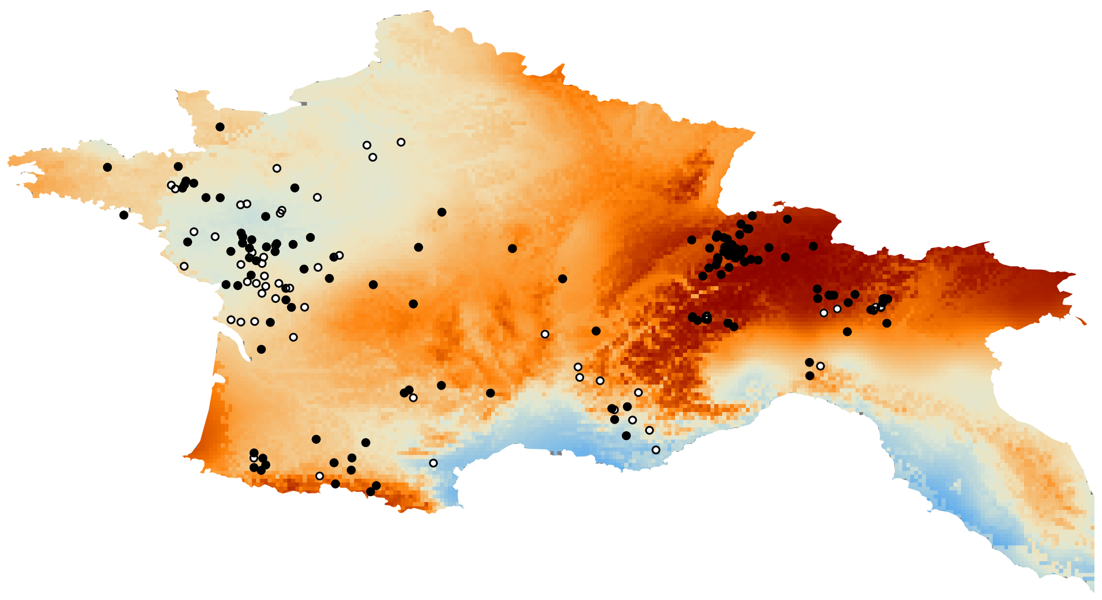

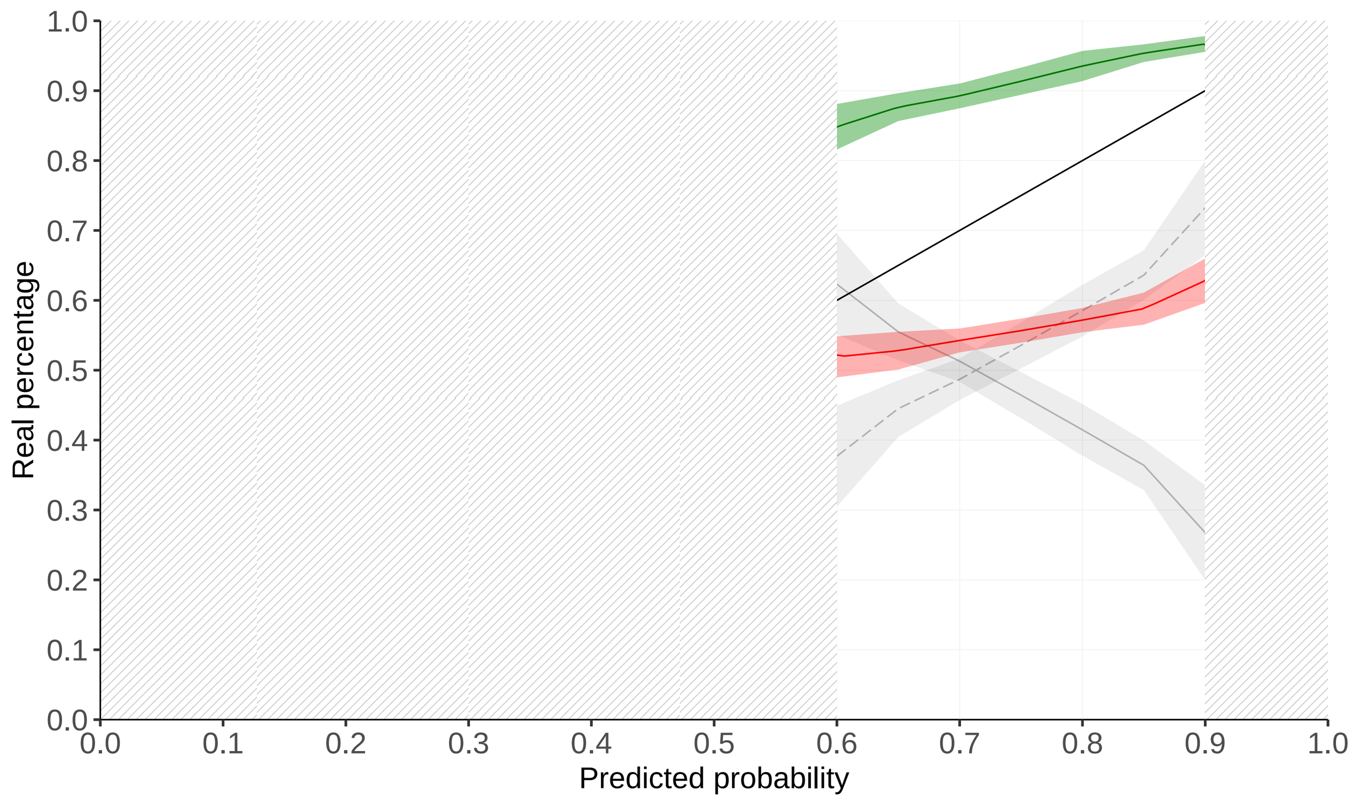


#### Chromosome 7: 39507171 CC Bio 18

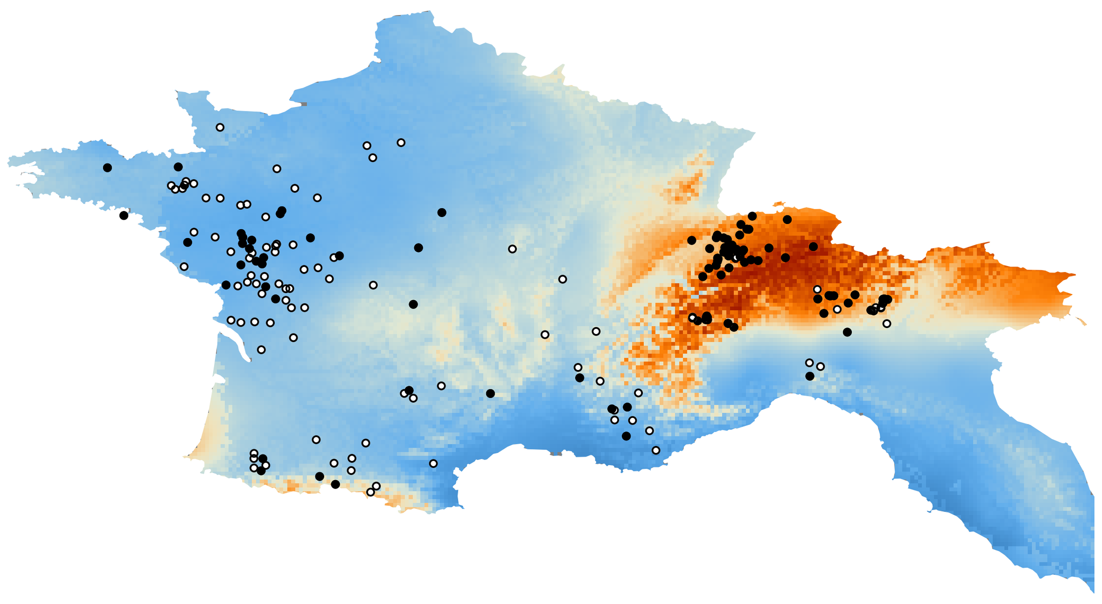

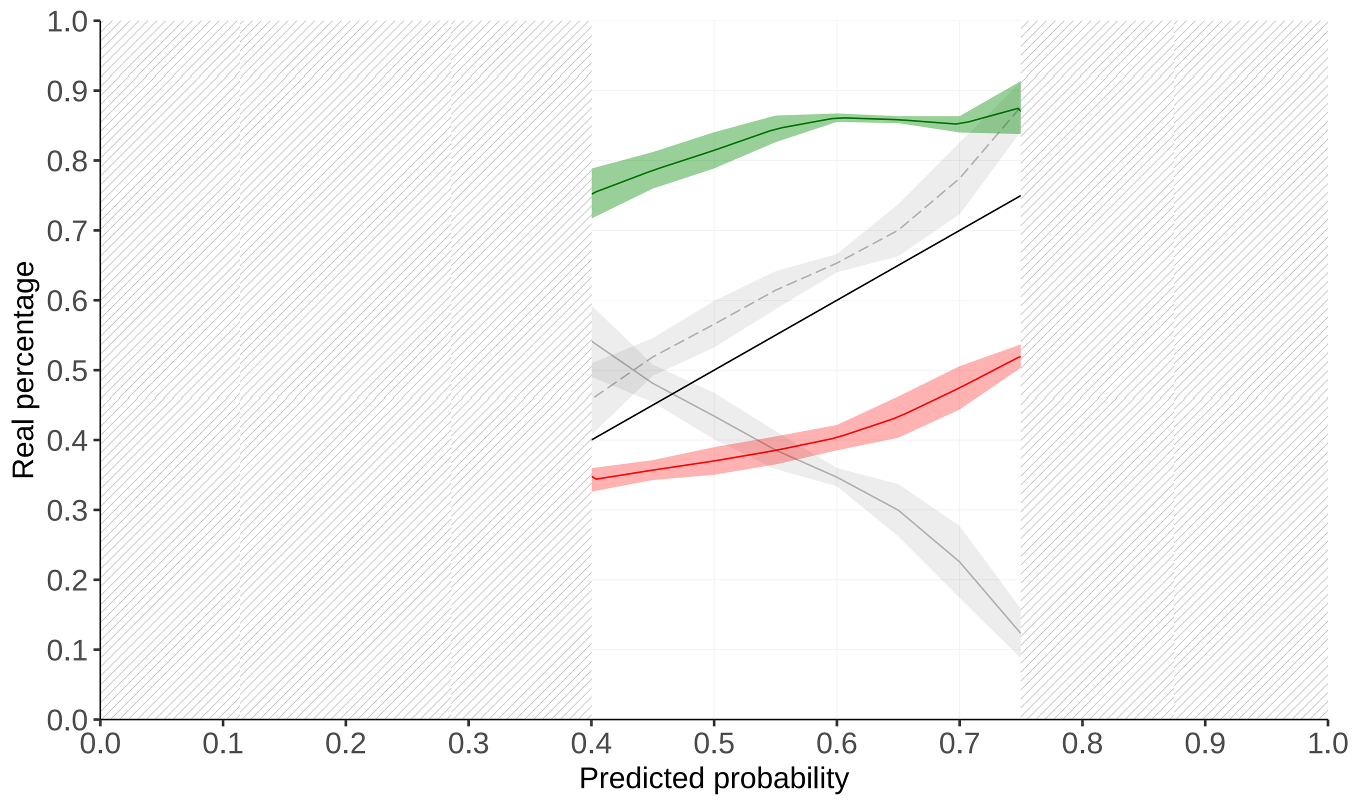


#### Chromosome 5: 6878482 GG Bio 3

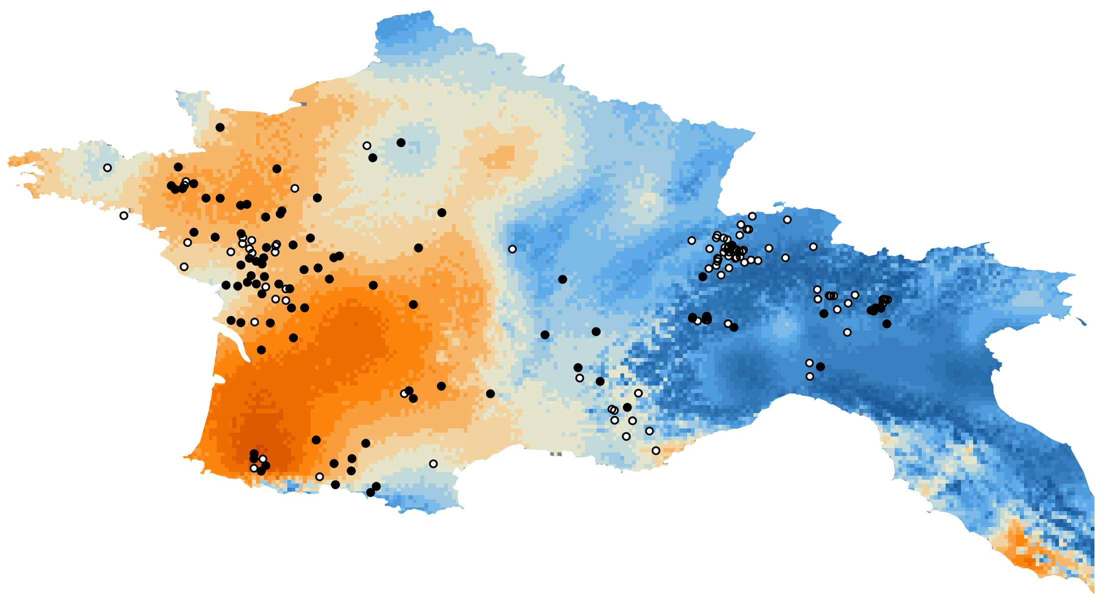

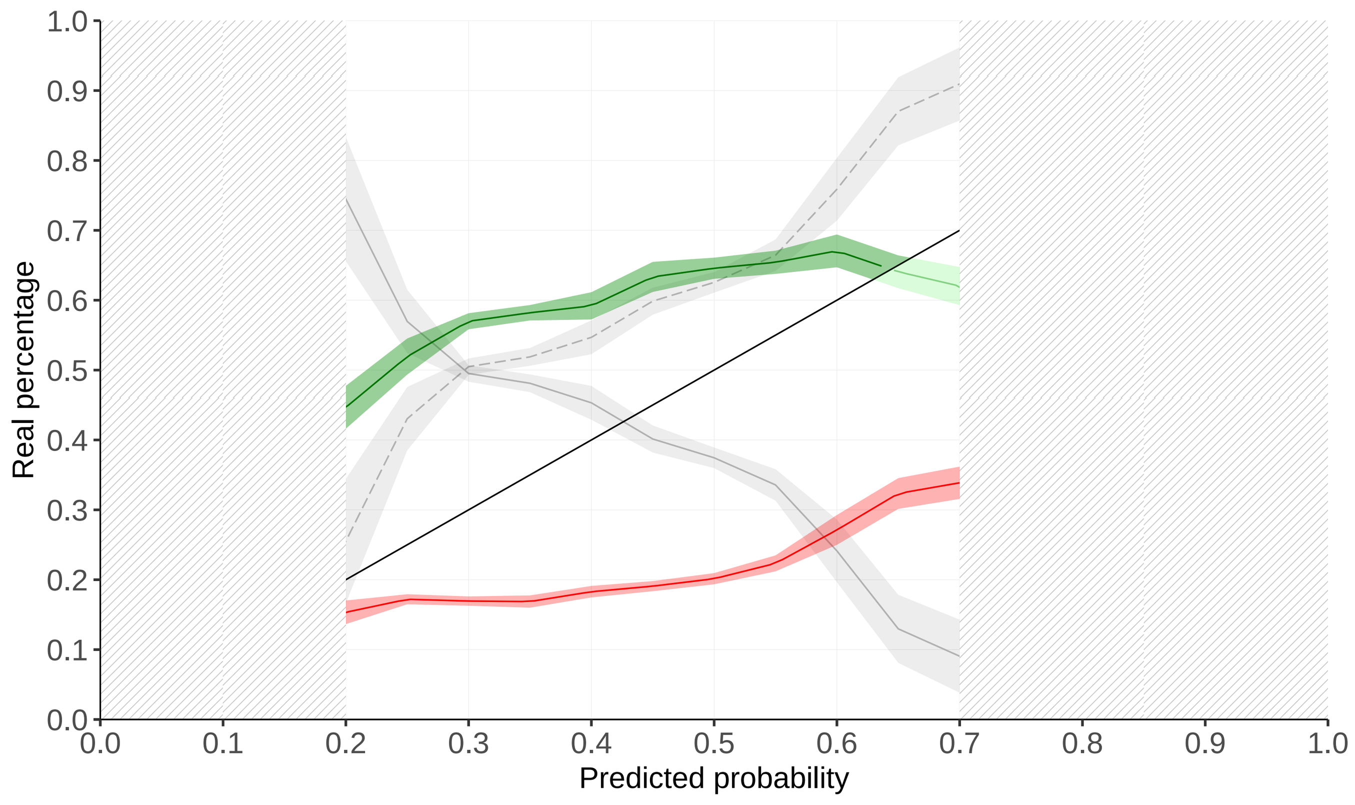


#### Chromosome 14: 8680762 AA Bio 18

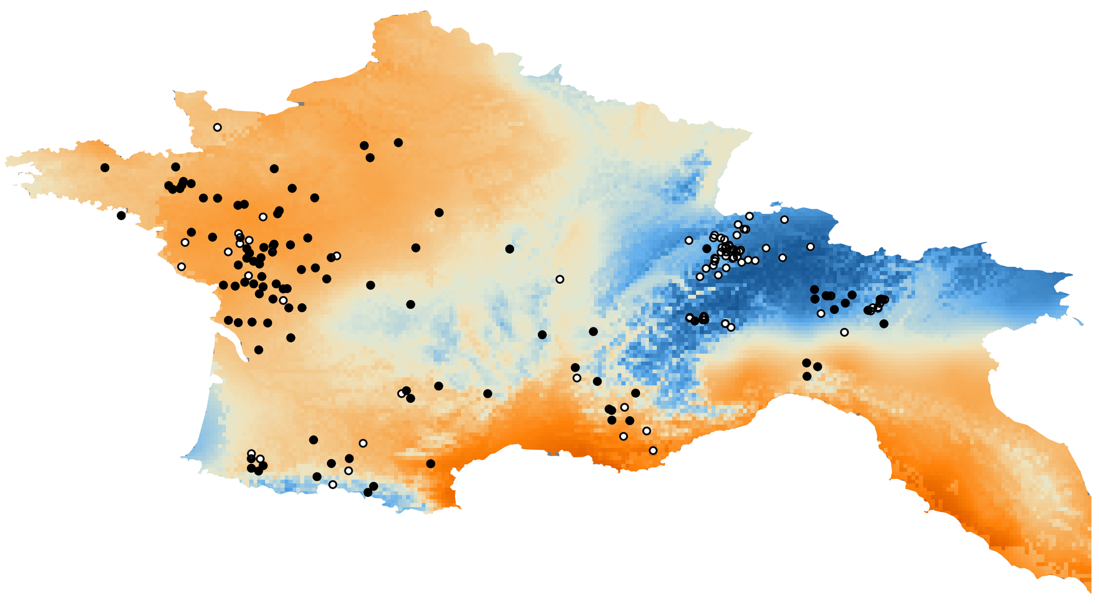

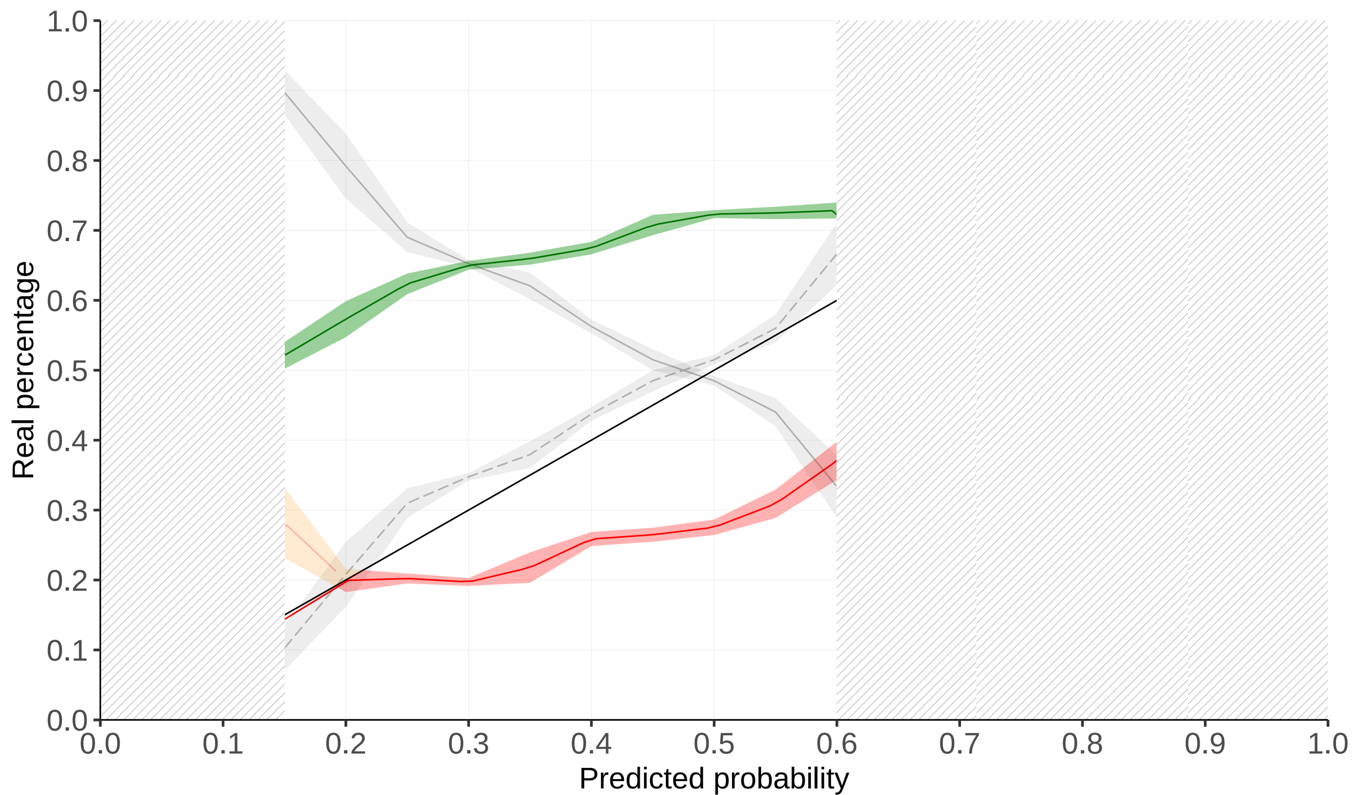
