## Supplementary material for "Spatial Areas of Genotype Probability (SPAG): predicting the spatial distribution of adaptive genetic variants under future climatic conditions": Supp. File 1-4

#### Suppl. File 1 - Method

##### Multivariate models

###### Intersection

From the theory of conditional probabilities, we know that the probability of the simultaneous presence of two genotypes G1 and G2 can be written:

$$p(G_1 \cap G_2) = p(G_1)\, p(G_2|G_1)$$

**Formula S1**

where $p(G1)$ is the probability of presence of the genotype G1, which can be computed using the univariate model (Formula 1 in the main text), and $p(G_2|G_1)$ is the conditional probability of G2 given G1. The computation of this second probability could be performed with a logistic regression where G1 is integrated as a covariate in the univariate model for G2 (Formula S2).

$$p(G_2|G_1) = \frac{e^{\beta_0+\beta_1 x_2+\beta_2 G_1}}{1+e^{\beta_0+\beta_1 x_2+\beta_2 G_1}}$$

**Formula S2**

where x2 is the environmental variable associated with G2 and $\beta_0$, $\beta_1$ and $\beta_2$ are the parameters of the logistic regression. However, as we would like to use this model to predict the probability of presence of the genotypes for any point of the region of interest, i.e also where values of G1 are not know, we suggest to estimate G1 by $p(G1)$ which can be computed for the entire territory from the univariate SPAG of G1. We therefore approximated $p(G_2|G_1)$ by $p(G_2|p(G_1))$ with the logistic regression in formula S3.

$$p(G_2|G_1) \approx \frac{e^{\beta_0+\beta_1 x_2+\beta_2 p(G_1)}}{1+e^{\beta_0+\beta_1 x_2+\beta_2 p(G_1)}}$$

**Formula S3**

The final SPAG $p(G_1 \cap G_2)$ can therefore be computed in three steps:

1. compute the SPAG corresponding to $p(G1)$ across the entire territory, using an univariate model with the environmental variable x1 associated with G1.

2. compute $p(G_2|G_1)$ using a logistic model with the G2 as the dependent variable, the environmental variable *x2* as the independent variable and the probability of presence *p(G1)* as a covariate (Formula S2).

3. multiply the results of $p(G1)$ obtained in step 1 with *p(G2|G1)* computed in step 2 to derive the final $p(G_1 \cap G_2)$.

Using the associative property of the intersection (i.e. $p(G_1 \cap G_2 \cap G_3) = p(G_3 \cap (G_1 \cap G_2))$), the procedure above can be extended to compute the probability of simultaneous presence of n genotypes of interest:

4. compute $p(G_3|(G_1 \cap G_2))$ using a logistic model with G3 as the dependent variable, the environmental variable x3 as the independent variable and the probability $p(G_1 \cap G_2)$ computed in step 3 as a covariate.

5. multiply the results of $p(G_1 \cap G_2)$ obtained in step 3 with $p(G_3|(G_1 \cap G_2))$ computed in step 4 to derive $p(G_1 \cap G_2 \cap G_3)$.

6. compute $p(G_4|(G_1 \cap G_2 \cap G_3))$ using a logistic model with G4 as the dependent variable, the environmental variable x4 as the independent variable and the probability $p(G_1 \cap G_2 \cap G_3)$ computed in step 5 as a covariate.

7. multiply the results of $p(G_1 \cap G_2 \cap G_3)$ obtained in step 5 with $p(G_1 \cap G_2 \cap G_3)$ computed in step 6 to derive $p(G_1 \cap G_2 \cap G_3 \cap G_4)$.

8. Carry on until obtaining $p(G_1 \cap G_2 \cap G_3 \cap G_4 \cap \ldots \cap G_n)$ for *n* genotypes of interest.

Note that this approach allows for the integration of adaptive genotypes associated with various environmental variables since the environmental variable *x1* used in step 1 to compute *p(G1)* can be different from the variable x2, x3, x4, etc. used in the following steps.

We implemented this recursive approach to build generalised intersection models predicting the simultaneous presence of *n* genotypes of interest as a R function *I-spag* available at : https://github.com/estellerochat/SPAG.

Union

To compute the probability of presence of at least one of the adaptive variant, we use the exclusion-inclusion principle, from which the probability of presence of genotypes *G1 OR G2* can be written:

$$p(G_1 \cup G_2) = p(G_1) + p(G_2) - p(G_1 \cap G_2)$$

**Formula S4**

where *p(G1)* and *p(G2)* can be computed with univariate SPAGs and *p(G1 ∩ G2)* using the intersection SPAG. For three genotypes, this becomes

$$p(G_1 \cup G_2 \cup G_3) = p(G_1) + p(G_2) + p(G_3) \\ -p(G_1 \cap G_2) - p(G_1 \cap G_3) - p(G_2 \cap G_3) \\ + p(G_1 \cap G_2 \cap G_3)$$

**Formula S5**

The generalisation for *n* genotypes is given by formula S6, and can be computed using the univariate and intersection models. This general case was implemented as an R function *U-spag* available at https://github.com/estellerochat/SPAG.

$$p\left(\bigcup_{i=1}^{n} G_i\right) = \sum_{i=1}^{n} p(G_i) - \sum_{i<j} p\left(G_i \bigcap G_j\right) + \sum_{i<j<k} p\left(G_i \bigcap G_j \bigcap G_k\right) + \ldots + (-1)^{n-1} p\left(\bigcap_{i=1}^{n} G_i\right)$$

$$=> p\left(\bigcup_{i=1}^{n} G_i\right) = \sum_{k=1}^{n}\left((-1)^{k-1} \sum_{1 \le i_1 < i_2 < \cdots < i_k < n} P(G_{i_1} \cap G_{i_2} \cap \ldots \cap G_{i_k})\right)$$

**Formula S6**

K-Percentage

To explain the K-Percentage method, we start with an example: if we have three genotypes, and would like to know the probability to carry at least 50% of them, this would be equivalent to the probability to carry at least two of the three genotypes. This can be expressed with Formula S7, which is the union of the probabilities of simultaneously carrying a combination of two genotypes chosen among the three.

$$p(50\%\, G_1, G_2, G_3) = p\big( (G_1 \cap G_2) \cup (G_1 \cap G_3) \cup (G_2 \cap G_3)\big)$$

**Formula S7**

By developing the union operators and summarising the results, we obtain:

$$p(50\%\, G_1, G_2, G_3) = p(G_1 \cap G_2) + p(G_1 \cap G_3) + p(G_2 \cap G_3) - 3p(G_1 \cap G_2 \cap G_3)$$

By generalizing this approach, the probability to carry at least K% of *n* adaptive variant can be written:

$$p(K\%\, G_{i=1\ldots.n}) = p\left(\bigcup_{i=1}^{n} \bigcap_{1 \le i_1 < i_2 < \cdots < i_{(K\% * n+1)}} (G_{i_1} \cap G_{i_2} \cap \ldots \cap G_{i_k})\right)$$

**Formula A8**

Again, this can be computed using univariate and intersection models and the general case was implemented as an R function *K-spag* available at https://github.com/estellerochat/SPAG.

#### Selection of training samples

In the cross-validation procedure, training individuals were not randomly selected, but were chosen such to represent the entire distribution of the environmental variable of interest in the study area. We thus retrieved the maximum and minimum value of the environmental variable at the sampled sites and divided this range into N uniform intervals, where N corresponds to the number of training individuals (25% of the total number of individuals). Since individuals are not necessarily sampled uniformly along the range of an environmental variable, we can obtain some intervals without any individuals and others with more than one individual. We therefore selected randomly one individual in each interval where individuals were present, and completed the training set with individuals randomly selected from all remaining individuals. We used this training set to calculate the SPAG.

#### Supp. File 2 – CDPOP simulation parameters

| | |
|---|---|
| looptime | 300 |
| cdclimgentime | 0 |
| matemoveno | 1 |
| matemoveparA | 0 |
| matemoveparB | 0 |
| matemoveparC | 0 |
| matemovethresh | 25max |
| sexans | Y |
| Freplace | Y |
| Mreplace | Y |
| philopatry | N |
| multiple_paternity | N |
| selfans | N |
| Fdispmoveno | 1 |
| FdispmoveparA | 0 |
| FdispmoveparB | 0 |
| FdispmoveparC | 0 |
| Fdispmovethresh | 25max |
| Mdispmoveno | 1 |
| MdispmoveparA | 0 |
| MdispmoveparB | 0 |
| MdispmoveparC | 0 |
| Mdispmovethresh | 25max |
| offno | 2 |

| | |
|---|---|
| Femalepercent | 50 |
| EqualsexratioBirth | N |
| TwinningPercent | 0 |
| popModel | exp |
| r | 1 |
| K_env | 0 |
| subpopmortperc | 0 |
| muterate | 0 |
| mutationtype | random |
| loci | 50 |
| intgenesans | random |
| allefreqfilename | N |
| alleles | 2 |
| mtdna | N |
| startGenes | 0 |
| cdevolveans | M_X3_L3_A2_ModelX |
| startSelection | 0 |
| betaFile_selection | see Figure 1 (main text) |
| epistasis | N |
| epigeneans | N |
| startEpigene | 0 |
| betaFile_epigene | N |
| cdinfect | N |
| transmissionprob | 0 |

### Supp. File 3 – Genetic data

A - Moroccan Dataset

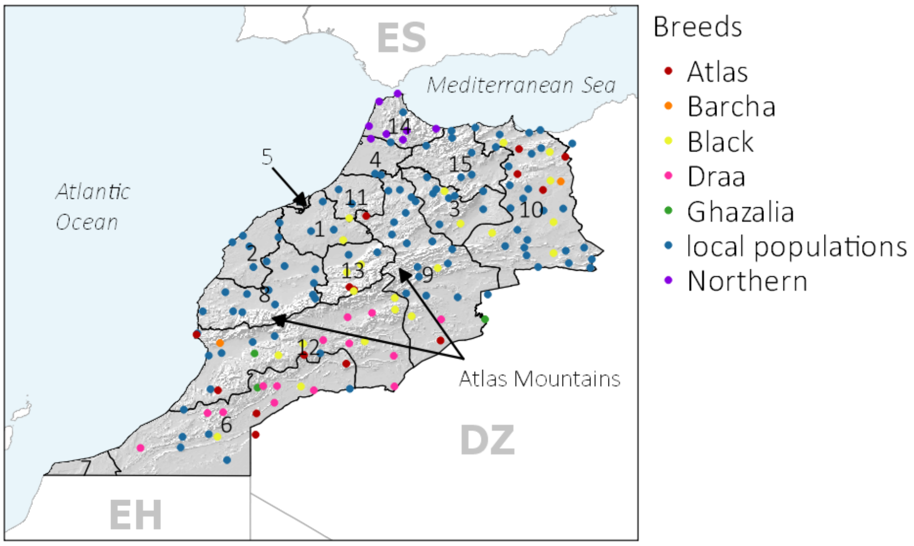


B - European Dataset

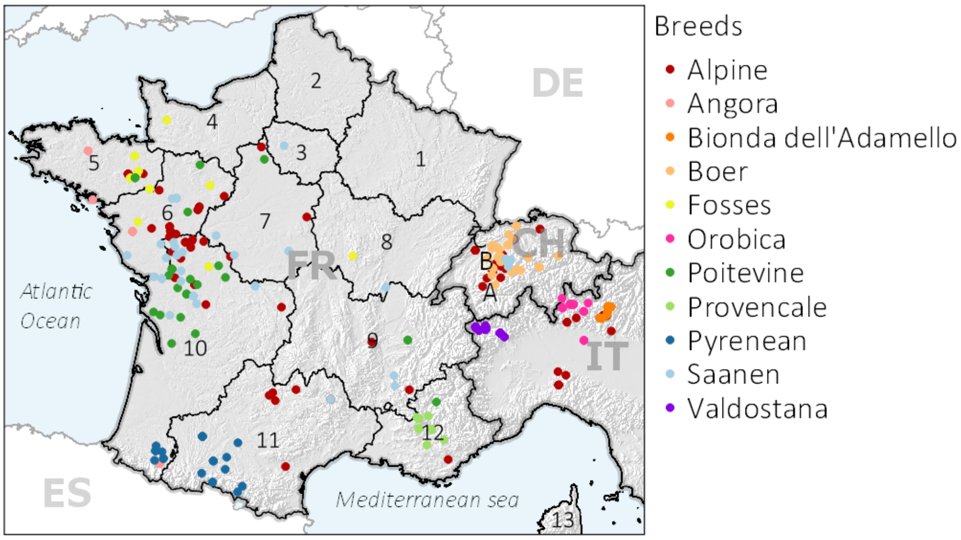

### <u>Supp. File 4 – Bioclimatic data</u>

| | |
|---|---|
| **BIO1** | Annual Mean Temperature |
| **BIO2** | Mean Diurnal Range (Mean of monthly (max temp - min temp)) |
| **BIO3** | Isothermality (BIO2/BIO7) (* 100) |
| **BIO4** | Temperature Seasonality (standard deviation *100) |
| **BIO5** | Max Temperature of Warmest Month |
| **BIO6** | Min Temperature of Coldest Month |
| **BIO7** | Temperature Annual Range (BIO5-BIO6) |
| **BIO8** | Mean Temperature of Wettest Quarter |
| **BIO9** | Mean Temperature of Driest Quarter |
| **BIO10** | Mean Temperature of Warmest Quarter |
| **BIO11** | Mean Temperature of Coldest Quarter |
| **BIO12** | Annual Precipitation |
| **BIO13** | Precipitation of Wettest Month |
| **BIO14** | Precipitation of Driest Month |
| **BIO15** | Precipitation Seasonality (Coefficient of Variation) |
| **BIO16** | Precipitation of Wettest Quarter |
| **BIO17** | Precipitation of Driest Quarter |
| **BIO18** | Precipitation of Warmest Quarter |
| **BIO19** | Precipitation of Coldest Quarter |
